# Genome-wide association analyses identify 44 risk variants and refine the genetic architecture of major depressive disorder

**DOI:** 10.1101/167577

**Authors:** Naomi R Wray, Stephan Ripke, Manuel Mattheisen, Maciej Trzaskowski, Enda M Byrne, Abdel Abdellaoui, Mark J Adams, Esben Agerbo, Tracy M Air, Till F M Andlauer, Silviu-Alin Bacanu, Marie Bækvad-Hansen, Aartjan T F Beekman, Tim B Bigdeli, Elisabeth B Binder, Douglas H R Blackwood, Julien Bryois, Henriette N Buttenschøn, Jonas Bybjerg-Grauholm, Na Cai, Enrique Castelao, Jane Hvarregaard Christensen, Toni-Kim Clarke, Jonathan R I Coleman, Lucía Colodro-Conde, Baptiste Couvy-Duchesne, Nick Craddock, Gregory E Crawford, Cheynna A Crowley, Hassan S Dashti, Gail Davies, Ian J Deary, Franziska Degenhardt, Eske M Derks, Nese Direk, Conor V Dolan, Erin C Dunn, Thalia C Eley, Nicholas Eriksson, Valentina Escott-Price, Farnush Farhadi Hassan Kiadeh, Hilary K Finucane, Andreas J Forstner, Josef Frank, Héléna A Gaspar, Michael Gill, Paola Giusti-Rorínguez, Fernando S Goes, Scott D Gordon, Jakob Grove, Lynsey S Hall, Christine Søholm Hansen, Thomas F Hansen, Stefan Herms, Ian B Hickie, Per Hoffmann, Georg Homuth, Carsten Horn, Jouke-Jan Hottenga, David M Hougaard, Ming Hu, Craig L Hyde, Marcus Ising, Rick Jansen, Fulai Jin, Eric Jorgenson, James A Knowles, Isaac S Kohane, Julia Kraft, Warren W. Kretzschmar, Jesper Krogh, Zoltan Kutalik, Jacqueline M Lane, Yihan Li, Yun Li, Penelope A Lind, Xiaoxiao Liu, Leina Lu, Donald J MacIntyre, Dean F MacKinnon, Robert M Maier, Wolfgang Maier, Jonathan Marchini, Hamdi Mbarek, Patrick McGrath, Peter McGuffin, Sarah E Medland, Divya Mehta, Christel M Middeldorp, Evelin Mihailov, Yuri Milaneschi, Lili Milani, Francis M Mondimore, Grant W Montgomery, Sara Mostafavi, Niamh Mullins, Matthias Nauck, Bernard Ng, Michel G Nivard, Dale R Nyholt, Paul F O’Reilly, Hogni Oskarsson, Michael J Owen, Jodie N Painter, Carsten Bøcker, Marianne Giørtz Pedersen, Roseann E. Peterson, Erik Pettersson, Wouter J Peyrot, Giorgio Pistis, Danielle Posthuma, Shaun M Purcell, Jorge A Quiroz, Per Qvist, John P Rice, Brien P. Riley, Margarita Rivera, Saira Saeed Mirza, Richa Saxena, Robert Schoevers, Eva C Schulte, Ling Shen, Jianxin Shi, Stanley I Shyn, Engilbert Sigurdsson, Grant C B Sinnamon, Johannes H Smit, Daniel J Smith, Hreinn Stefansson, Stacy Steinberg, Craig A Stockmeier, Fabian Streit, Jana Strohmaier, Katherine E Tansey, Henning Teismann, Alexander Teumer, Wesley Thompson, Pippa a Thomson, Thorgeir E Thorgeirsson, Chao Tian, Matthew Traylor, Jens Treutlein, Vassily Trubetskoy, André G Uitterlinden, Daniel Umbricht, Sandra Van der Auwera, Albert M van Hemert, Alexander Viktorin, Peter M Visscher, Yunpeng Wang, Bradley T. Webb, Shantel Marie Weinsheimer, Jürgen Wellmann, Gonneke Willemsen, Stephanie H Witt, Yang Wu, Hualin S Xi, Jian Yang, Futao Zhang, eQTLGen Consortium, 23andMe Research Team, Volker Arolt, Bernhard T Baune, Klaus Berger, Dorret I Boomsma, Sven Cichon, udo Dannlowski, EJC de Geus, J Raymond DePaulo, Enrico Domenici, Katharina Domschke, Tönu Esko, Hans J Grabe, Steven P Hamilton, Caroline Hayward, Andrew C Heath, David A Hinds, Kenneth S Kendler, Stefan Kloiber, Glyn Lewis, Qingqin S Li, Susanne Lucae, Pamela AF Madden, Patrik K Magnusson, Nicholas G Martin, Andrew M McIntosh, Andres Metspalu, Ole Mors, Preben Bo Mortensen, Bertram Müller-Myhsok, Merete Nordentoft, Markus M Nöthen, Michael C O’Donovan, Sara A Paciga, Nancy L Pedersen, Brenda WJH Penninx, Roy H Perlis, David J Porteous, James B Potash, Martin Preisig, Marcella Rietschel, Catherine Schaefer, Thomas G Schulze, Jordan W Smoller, Kari Stefansson, Henning Tiemeier, Rudolf Uher, Henry Völzke, Myrna M Weissman, Thomas Werge, Ashley R Winslow, Cathryn M Lewis, Douglas F Levinson, Gerome Breen, Anders D Børglum, Patrick F Sullivan, for the Major Depressive Disorder Working Group of the Psychiatric Genomics Consortium

## Abstract

Major depressive disorder (MDD) is a notably complex illness with a lifetime prevalence of 14%.^1^ It is often chronic or recurrent and is thus accompanied by considerable morbidity, excess mortality, substantial costs, and heightened risk of suicide.^2-7^ MDD is a major cause of disability worldwide.^8^ We conducted a genome-wide association (GWA) meta-analysis in 130,664 MDD cases and 330,470 controls, and identified 44 independent loci that met criteria for statistical significance. We present extensive analyses of these results which provide new insights into the nature of MDD. The genetic findings were associated with clinical features of MDD, and implicated prefrontal and anterior cingulate cortex in the pathophysiology of MDD (regions exhibiting anatomical differences between MDD cases and controls). Genes that are targets of antidepressant medications were strongly enriched for MDD association signals (P=8.5×10^−10^), suggesting the relevance of these findings for improved pharmacotherapy of MDD. Sets of genes involved in gene splicing and in creating isoforms were also enriched for smaller MDD GWA P-values, and these gene sets have also been implicated in schizophrenia and autism. Genetic risk for MDD was correlated with that for many adult and childhood onset psychiatric disorders. Our analyses suggested important relations of genetic risk for MDD with educational attainment, body mass, and schizophrenia: the genetic basis of lower educational attainment and higher body mass were putatively causal for MDD whereas MDD and schizophrenia reflected a partly shared biological etiology. All humans carry lesser or greater numbers of genetic risk factors for MDD, and a continuous measure of risk underlies the observed clinical phenotype. MDD is not a distinct entity that neatly demarcates normalcy from pathology but rather a useful clinical construct associated with a range of adverse outcomes and the end result of a complex process of intertwined genetic and environmental effects. These findings help refine and define the fundamental basis of MDD.

---

Twin studies attribute ~40% of the variation in liability to MDD to additive genetic effects (heritability, *h*^2^),^9^ and *h*^2^ may be greater for recurrent, early-onset, and postpartum MDD.^10,11^ GWA studies of MDD have had notable difficulties in identifying loci.^12^ Previous findings suggest that an appropriately designed study should identify susceptibility loci. Direct estimates of the proportion of variance attributable to genome-wide SNPs (SNP heritability, 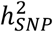 indicate that around a quarter of the *h*^2^ for MDD is due to common genetic variants.^13,14^ Although there were no significant findings in the initial Psychiatric Genomics Consortium (PGC) MDD mega-analysis (9,240 MDD cases)^15^ or in the CHARGE meta-analysis of depressive symptoms (34,549 respondents),^16^ more recent studies have proven modestly successful. A study of Han Chinese women (5,303 MDD cases) identified two genome-wide significant loci,^17^ a meta-analysis of depressive symptoms (161,460 individuals) identified two loci,^18^ and an analysis of self-reported MDD identified 15 loci (75,607 cases).^19^

There are many reasons why identifying causal loci for MDD has proven difficult.^12^ MDD is probably influenced by many genetic loci each with small effects,^20^ as are most common complex human diseases^21^ including psychiatric disorders.^22,23^ A major lesson in human complex trait genetics is that large samples are essential, especially for common and etiologically heterogeneous illnesses like MDD.^24^ We sought to accumulate a large sample to identify common genetic variation involved in the etiology of MDD.^24^

## Analysis of MDD anchor with six expanded cohorts shows polygenic prediction & clinical relevance

We defined an “anchor” cohort of 29 samples that mostly applied standard methods for assessing MDD (***Table S1***). MDD cases in the anchor cohort were traditionally ascertained and typically characterized (i.e., using direct interviews with structured diagnostic instruments). We identified six “expanded” cohorts that used alternative methods to identify MDD (***Table S2***; deCODE, Generation Scotland, GERA, iPSYCH, UK Biobank, and 23andMe, Inc.). All seven cohorts focused on clinically-significant MDD. We evaluated the comparability of these cohorts (***Table S3***) by estimating the common-variant genetic correlations (*r_g_*) of the anchor cohort with the expanded cohorts. These analyses strongly supported the comparability of the seven cohorts (***Table S4***) as the weighted mean *r_g_* was 0.76 (SE 0.028) with no statistical evidence of heterogeneity in the *r_g_* estimates (*P*=0.13). As a benchmark for the MDD *r_g_* estimates, the weighted mean *r_g_* between schizophrenia cohorts was 0.84 (SE 0.05).^13^

We completed a GWA meta-analysis of 9.6 million imputed SNPs in seven cohorts containing 130,664 MDD cases and 330,470 controls (***Figure 1***; full details in ***Online Methods***). There was no evidence of uncontrolled inflation (LD score regression intercept 1.018, SE 0.009). We estimated 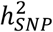 to be 8.9% (SE 0.004, liability scale, assuming lifetime population risk of 0.15), and this is around a quarter of *h*^2^ estimated from twin or family studies.^9^ This fraction is somewhat lower than that of other complex traits,^21^ and is plausibly due to etiological heterogeneity.

**Figure 1:**
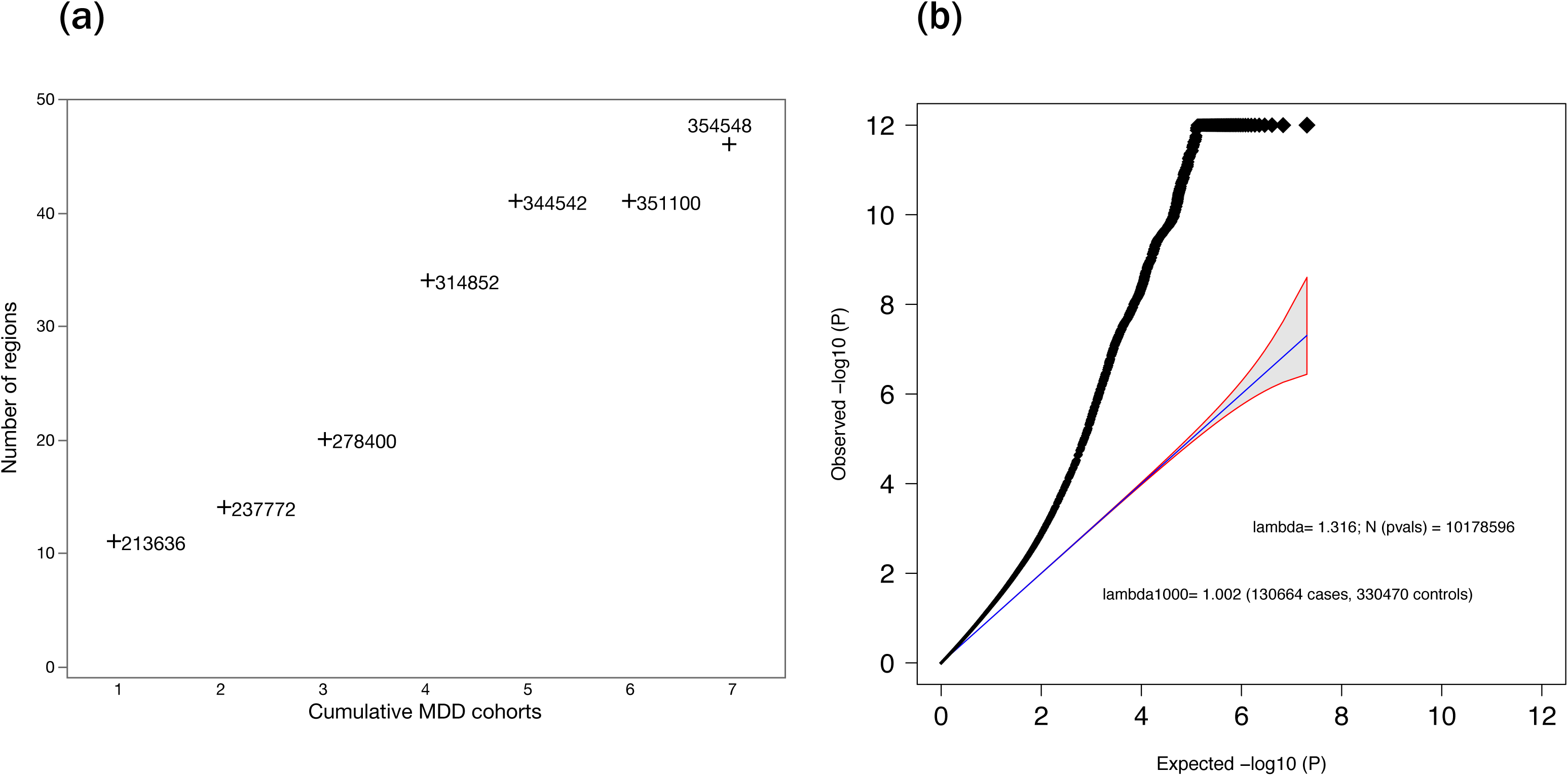
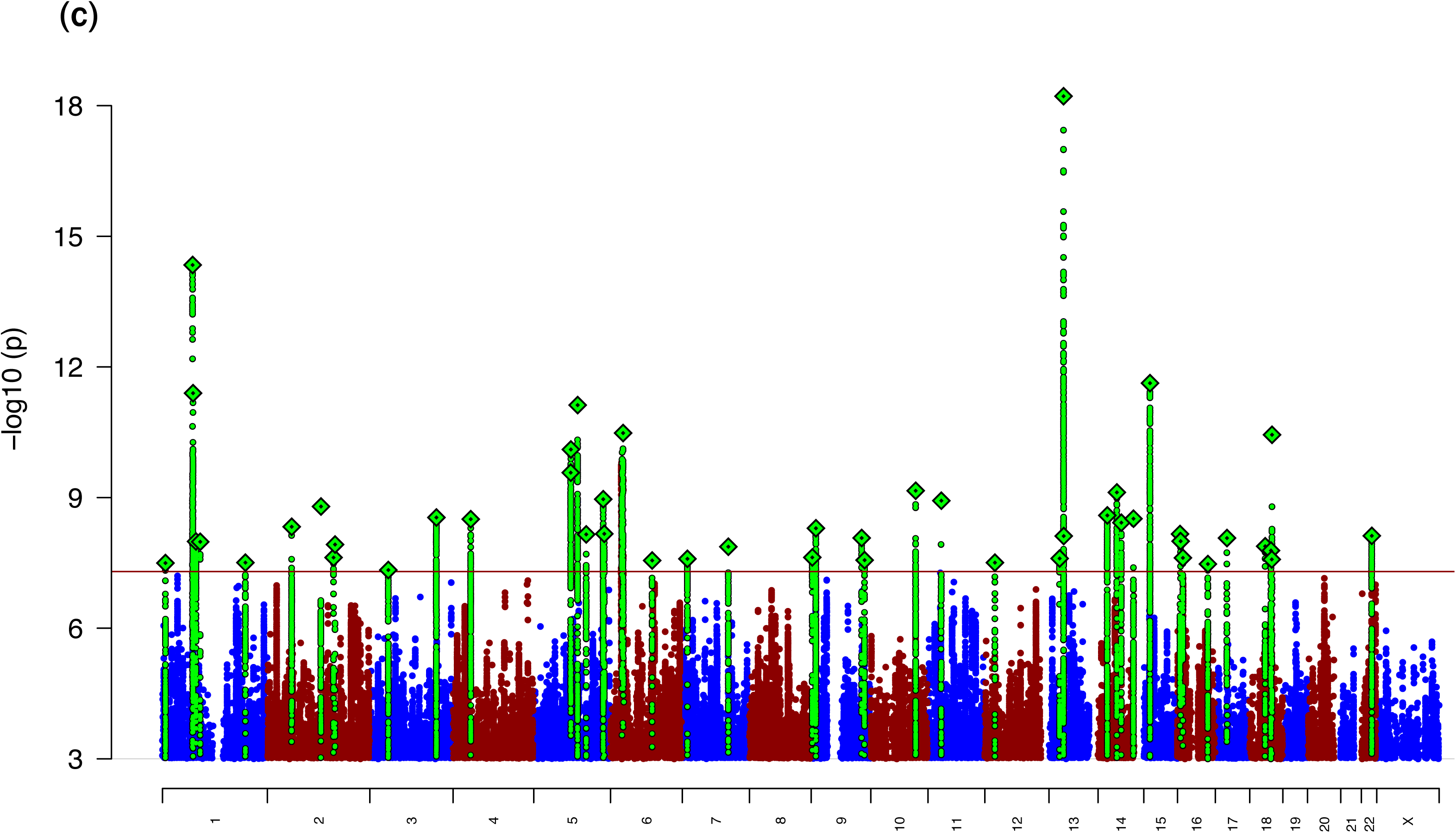
Results of GWA meta-analysis of seven cohorts for MDD. (a) Relation between adding cohorts and number of genome-wide significant genomic regions. Beginning with the largest cohort (1), added the next largest cohort (2) until all cohorts were included (7). The number next to each point shows the total effective sample size. (b) Quantile-quantile plot showing a marked departure from a null model of no associations (the y-axis is truncated at 1e-12). (c) Manhattan plot with x-axis showing genomic position (chr1-chr22), and the y-axis showing statistical significance as −log_10_(P). The red line shows the genome-wide significance threshold (P=5×10^8^).

We used genetic risk score (GRS) analyses to demonstrate the validity of our GWA results for clinical MDD (***Figure 2***). As expected, the variance explained in out-of-sample prediction increased with the size of the GWA discovery cohort (***Figure 2a***). Across all samples in the anchor cohort, GRS explained 1.9% of variance in liability (***Figure S1a***), GRS ranked cases higher than controls with probability 0.57, and the odds ratio of MDD for those in the 10^th^ versus 1^st^ GRS decile (OR10) was 2.4 (***Figure 2b***, ***Table S5***). GRS were significantly higher in those with more severe MDD, as measured in different ways (***Figure 2c***)

**Figure 2:**
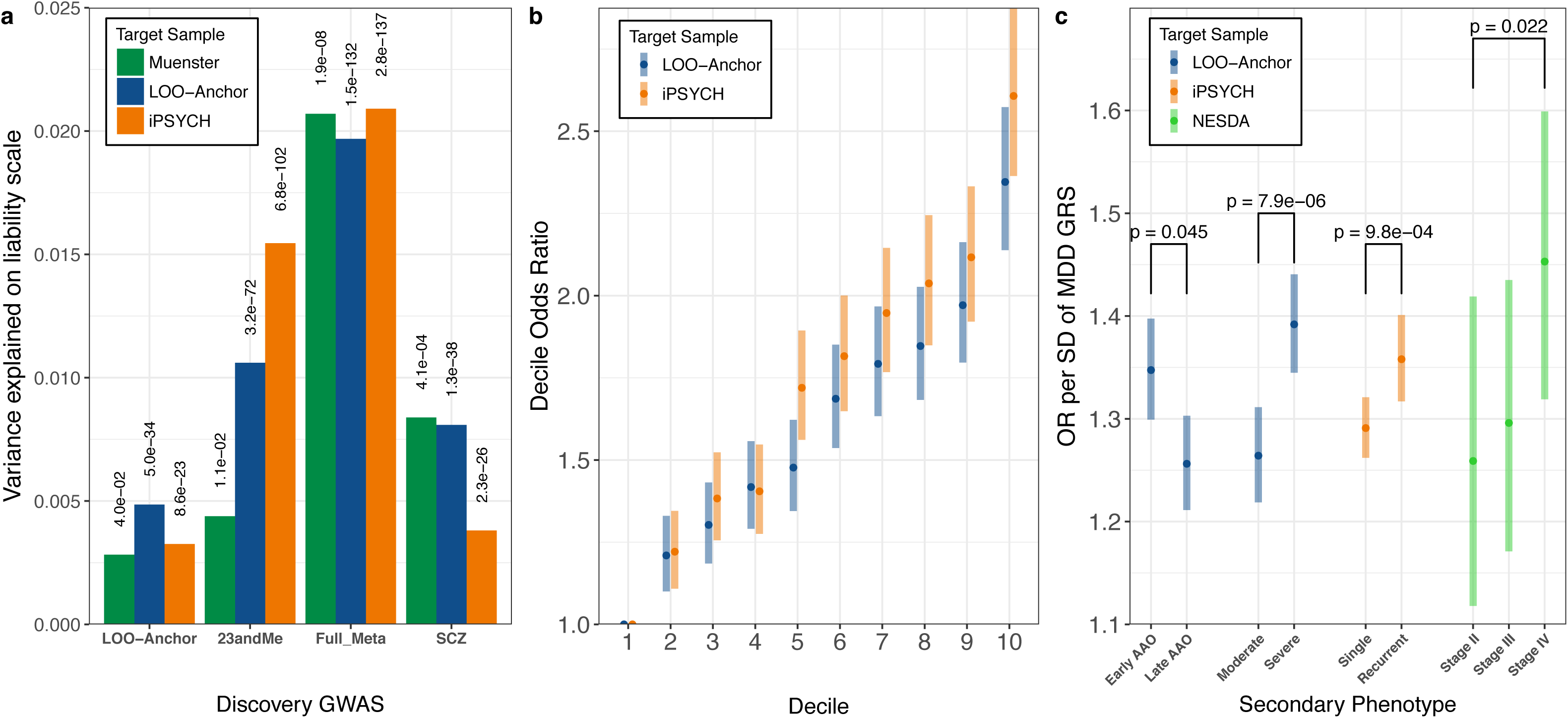
Out-of-sample genetic risk score (GRS) prediction analyses. (a) Variance explained on the liability scale based on different discovery samples for three target samples: anchor cohort (16,823 cases, 25,632 controls), iPSYCH (a nationally representative sample of 18,629 cases and 17,841 controls) and a clinical cohort from Münster not included in the GWA analysis (845 MDD inpatient cases, 834 controls). The anchor cohort is included as both discovery and target as we computed out-of-sample GRS for each anchor cohort sample, combined the results, and modeled case-control status as predicted by standardized GRS and cohort (see **Online Methods**). (b) Odd ratios of MDD per GRS decile relative to the first decile for iPSYCH and anchor cohorts. (c) MDD GRS (from out-of-sample discovery sets) were significantly higher in MDD cases with: earlier age at onset; more severe MDD symptoms (based on number of criteria endorsed); recurrent MDD compared to single episode; and chronic/unremitting MDD (“Stage IV” compared to “Stage II”, first-episode MDD^103^). Error bars represent 95% confidence intervals.

## Implications of the individual loci for the biology of MDD

Our meta-analysis of seven MDD cohorts identified 44 independent loci that were statistically significant (*P*<5×10^−8^), statistically independent of any other signal,^25^ supported by multiple SNPs, and showed consistent effects across cohorts. This number is consistent with our prediction that MDD GWA discovery would require about five times more cases than for schizophrenia (lifetime risk ~1% and *h*^2^~0.8) to achieve approximately similar power.^26^ Of these 44 loci, 30 are novel and 14 were significant in a prior study of MDD or depressive symptoms (the overlap of our findings: 1/1 with the CHARGE depressive symptom study,^16^ 0/2 overlap with CONVERGE MDD study,^17^ 1/2 overlap with the SSGAC depressive symptom study,^18^ and 13/16 overlap with 23andMe self-report of MDD^19^). There are few trans-ancestry comparisons for MDD so we contrasted these European results with the Han Chinese CONVERGE study (***Online Methods***).

***Table 1*** lists genes in or near the lead SNP in each region, regional plots are in the *Supplemental File*, and ***Table S6*** provides extensive summaries of available information about the biological functions of the genes in each region. In nine of the 44 loci, the lead SNP is within a gene, there is no other gene within 200 kb, and the gene is known to play a role in neuronal development, synaptic function, transmembrane adhesion complexes, and/or regulation of gene expression in brain.

**Table 1.** Genomic regions significantly associated with MDD

| Chr | Region (Mb) | SNP | Location-bp | P | A1/2 | OR-A1 (SE) | Frq | Prev | Gene Context |
| --- | --- | --- | --- | --- | --- | --- | --- | --- | --- |
| 1 | 8.390-8.895 | rs159963 | 8,504,421 | 3.2E-08 | A/C | 0.97 (0.0049) | 0.56 | H,S | [RERE]; SLC45A1,100194 |
| 1 | 72.511-73.059 | rs1432639 | 72,813,218 | 4.6E-15 | A/C | 1.04 (0.005) | 0.63 | H | NEGR1,-64941 |
| 1 | 73.275-74.077 | rs12129573 | 73,768,366 | 4.0E-12 | A/C | 1.04 (0.005) | 0.37 | H,S | LINC01360,-3486 |
| 1 | 80.785-80.980 | rs2389016 | 80,799,329 | 1.0E-08 | T/C | 1.03 (0.0053) | 0.28 |  |  |
| 1 | 90.671-90.966 | rs4261101 | 90,796,053 | 1.0E-08 | A/G | 0.97 (0.005) | 0.37 |  |  |
| 1 | 197.343-197.864 | rs9427672 | 197,754,741 | 3.1E-08 | A/G | 0.97 (0.0058) | 0.24 |  | DENND1B,-10118 |
| 2 | 57.765-58.485 | rs11682175 | 57,987,593 | 4.7E-09 | T/C | 0.97 (0.0048) | 0.52 | H,S | VRK2,-147192 |
| 2 | 156.978-157.464 | rs1226412 | 157,111,313 | 2.4E-08 | T/C | 1.03 (0.0059) | 0.79 |  | [LINC01876]; NR4A2,69630; GPD2,-180651 |
| 3 | 44.222-44.997 | chr3_44287760_I | 44,287,760 | 4.6E-08 | I/D | 1.03 (0.0051) | 0.34 | T | [TOPAZ1]; TCAIM,-91850; ZNF445,193501 |
| 3 | 157.616-158.354 | rs7430565 | 158,107,180 | 2.9E-09 | A/G | 0.97 (0.0048) | 0.58 | H | [RSRC1]; LOC100996447,155828; MLF1,-181772 |
| 4 | 41.880-42.189 | rs34215985 | 42,047,778 | 3.1E-09 | C/G | 0.96 (0.0063) | 0.24 |  | [SLC30A9]; LINC00682,-163150; DCAF4L1,59294 |
| 5 | 87.443-88.244 | chr5_87992715_I | 87,992,715 | 7.9E-11 | I/D | 0.97 (0.005) | 0.58 | H | LINC00461,-12095; MEF2C,21342 |
| 5 | 103.672-104.092 | chr5_103942055_D | 103,942,055 | 7.5E-12 | I/D | 1.03 (0.0048) | 0.48 | C |  |
| 5 | 124.204-124.328 | rs116755193 | 124,251,883 | 7.0E-09 | T/C | 0.97 (0.005) | 0.38 |  | LOC101927421,-120640 |
| 5 | 164.440-164.789 | rs11135349 | 164,523,472 | 1.1E-09 | A/C | 0.97 (0.0048) | 0.48 | H |  |
| 5 | 166.977-167.056 | rs4869056 | 166,992,078 | 6.8E-09 | A/G | 0.97 (0.005) | 0.63 |  | [TENM2] |
| 6 | 27.738-32.848 | rs115507122 | 30,737,591 | 3.3E-11 | C/G | 0.96 (0.0063) | 0.18 | S | extended MHC |
| 6 | 99.335-99.662 | rs9402472 | 99,566,521 | 2.8E-08 | A/G | 1.03 (0.0059) | 0.24 |  | FBXL4,-170672; C6orf168,154271 |
| 7 | 12.154-12.381 | rs10950398 | 12,264,871 | 2.6E-08 | A/G | 1.03 (0.0049) | 0.41 |  | [TMEM106B]; VWDE,105637 |
| 7 | 108.925-109.230 | rs12666117 | 109,105,611 | 1.4E-08 | A/G | 1.03 (0.0048) | 0.47 |  |  |
| 9 | 2.919-3.009 | rs1354115 | 2,983,774 | 2.4E-08 | A/C | 1.03 (0.0049) | 0.62 | H | PUM3,-139644; LINC01231,-197814 |
| 9 | 11.067-11.847 | rs10959913 | 11,544,964 | 5.1E-09 | T/G | 1.03 (0.0057) | 0.76 |  |  |
| 9 | 119.675-119.767 | rs7856424 | 119,733,595 | 8.5E-09 | T/C | 0.97 (0.0053) | 0.29 |  | [ASTN2] |
| 9 | 126.292-126.735 | rs7029033 | 126,682,068 | 2.7E-08 | T/C | 1.05 (0.0093) | 0.07 |  | [DENND1A]; LHX2,-91820 |
| 10 | 106.397-106.904 | rs61867293 | 106,563,924 | 7.0E-10 | T/C | 0.96 (0.0061) | 0.20 | H | [SORCS3] |
| 11 | 31.121-31.859 | rs1806153 | 31,850,105 | 1.2E-09 | T/G | 1.04 (0.0059) | 0.22 |  | [DKFZp686K1684]; [PAUPAR]; ELP4,44032; |
| 12 | 23.924-24.052 | rs4074723 | 23,947,737 | 3.1E-08 | A/C | 0.97 (0.0049) | 0.41 |  | [SOX5] |
| 13 | 44.237-44.545 | rs4143229 | 44,327,799 | 2.5E-08 | A/C | 0.95 (0.0091) | 0.92 |  | [ENOX1]; LACC1,-125620; CCDC122,82689 |
| 13 | 53.605-54.057 | rs12552 | 53,625,781 | 6.1E-19 | A/G | 1.04 (0.0048) | 0.44 | H | [OLFM4]; LINC01065,80099 |
| 14 | 41.941-42.320 | rs4904738 | 42,179,732 | 2.6E-09 | T/C | 0.97 (0.0049) | 0.57 |  | [LRFN5] |
| 14 | 64.613-64.878 | rs915057 | 64,686,207 | 7.6E-10 | A/G | 0.97 (0.0049) | 0.42 |  | [SYNE2]; MIR548H1,-124364; ESR2,7222 |
| 14 | 75.063-75.398 | chr14_75356855_I | 75,356,855 | 3.8E-09 | D/I | 1.03 (0.0049) | 0.49 |  | [DLST]; PROX2,-26318; RPS6KL1,13801 |
| 14 | 103.828-104.174 | rs10149470 | 104,017,953 | 3.1E-09 | A/G | 0.97 (0.0049) | 0.49 | S | BAG5,4927; APOPT1,-11340 |
| 15 | 37.562-37.929 | rs8025231 | 37,648,402 | 2.4E-12 | A/C | 0.97 (0.0048) | 0.57 | H |  |
| 16 | 6.288-6.347 | rs8063603 | 6,310,645 | 6.9E-09 | A/G | 0.97 (0.0053) | 0.65 |  | [RBFOX1] |
| 16 | 7.642-7.676 | rs7198928 | 7,666,402 | 1.0E-08 | T/C | 1.03 (0.005) | 0.62 |  | [RBFOX1] |
| 16 | 13.022-13.119 | rs7200826 | 13,066,833 | 2.4E-08 | T/C | 1.03 (0.0055) | 0.25 |  | [SHISA9]; CPPED1,-169089 |
| 16 | 71.631-72.849 | rs11643192 | 72,214,276 | 3.4E-08 | A/C | 1.03 (0.0049) | 0.41 |  | PMFBP1,-7927; DHX38,67465; |
| 17 | 27.345-28.419 | rs17727765 | 27,576,962 | 8.5E-09 | T/C | 0.95 (0.0088) | 0.92 |  | [CRYBA1]; MYO18A,-69555; NUFIP2,5891 |
| 18 | 36.588-36.976 | rs62099069 | 36,883,737 | 1.3E-08 | A/T | 0.97 (0.0049) | 0.42 |  | [MIR924HG] |
| 18 | 50.358-50.958 | rs11663393 | 50,614,732 | 1.6E-08 | A/G | 1.03 (0.0049) | 0.45 | O | [DCC]; MIR4528,-148738 |
| 18 | 51.973-52.552 | rs1833288 | 52,517,906 | 2.6E-08 | A/G | 1.03 (0.0054) | 0.72 |  | [RAB27B]; CCDC68,50833 |
| 18 | 52.860-53.268 | rs12958048 | 53,101,598 | 3.6E-11 | A/G | 1.03 (0.0051) | 0.33 | S | [TCF4]; MIR4529,-44853 |
| 22 | 40.818-42.216 | rs5758265 | 41,617,897 | 7.6E-09 | A/G | 1.03 (0.0054) | 0.28 | H,S | [L3MBTL2]; EP300-AS1,-24392; CHADL,7616 |

Shown are data for 44 genomic loci achieving genome-wide significance for MDD. Chr (chromosome) and Region (boundaries in Mb, hg19) are shown, defined by locations of SNPs with $P < 1 \times 10^{-5}$ and $LD r^2 > 0.1$ with the most associated SNP (lowest P-value in region, listed) whose location is given in bp. In three regions a second SNP fulfils the filtering criteria and these were followed up with conditional analyses: Chr1: conditional analysis selects only rs1432639 as significant, with $P = 2.0 \times 10^{-4}$ for rs12134600 after fitting rs1432639; Chr5, conditional analysis shows two independent associations selecting rs247910 and rs10514301 as the most associated SNPs; and Chr10 conditional analysis selects only rs61867293 with $P = 8.6 \times 10^{-5}$ for rs1021363 after conditioning on rs61867293. For each of the 47 SNPs, there is at least 1 additional genome-wide significant SNP in the cluster of surrounding SNPs with low P-values. Chromosome X was analyzed but had no findings that met genome-wide significance.
Column labels and abbreviations. A1/2 = the two alleles (or insertion-deletion); A1 was tested for association, and its OR (odds ratio) and SE (standard error) are shown. FreqU = frequency of A1 in controls across all cohorts. Entries in the “Prev” column indicate which of four previous studies identified genome-significant associations in a region. H=Hyde et al.,<sup>19</sup> 23andMe GWA of self-reported clinical depression (discovery sample overlaps with this paper); O=Okbay et al.,<sup>18</sup> meta-analysis of GWA of MDD, depressive symptoms, psychological well-being and neuroticism (includes many anchor cohorts from this paper); S=PGC report on 108 schizophrenia-associated loci<sup>22</sup>; and C=CHARGE consortium meta-analysis of depressive symptoms in primarily older adults.<sup>16</sup> Gene context: distances between the Peak SNP and the closest genes are shown. Brackets indicate that the Peak SNP was within that gene. The closest genes upstream (taking strand into account, as a negative number indicating distance in bp between Peak SNP and the nearest gene boundary) and downstream (positive distance in bp) are also shown, if there is a flanking gene within 200 kb. The name of the closest gene is bolded. Note that it is generally not known whether the associated SNPs have biological effects on these or other more distant genes.

The two most significant SNPs are located in or near *OLFM4* and *NEGR1*, which were previously associated with obesity and body mass index.^27-32^ *OLFM4* (olfactomedin 4) has diverse functions outside the CNS including myeloid precursor cell differentiation, innate immunity, anti-apoptotic effects, gut inflammation, and is over-expressed in diverse common cancers.^33^ Many olfactomedins also have roles in neurodevelopment and synaptic function;^34^ e.g., latrophilins form trans-cellular complexes with neurexins^35^ and with *FLRT3* to regulate glutamatergic synapse number.^36^ *Olfm4* was highly upregulated after spinal transection, possibly related to inhibition of subsequent neurite outgrowth.^37^ *NEGR1* (neuronal growth regulator 1) influences axon extension and synaptic plasticity in cortex, hypothalamus, and hippocampus,^38-40^ and modulates synapse formation in hippocampus^41,42^ via regulation of neurite outgrowth.^43,44^ High expression, modulated by nutritional state, is seen in brain areas relevant to feeding, suggesting a role in control of energy intake.^45^ The same SNP alleles are associated with increased risk of obesity and MDD (see also Mendelian randomization analyses below) and are associated with *NEGR1* gene expression in brain (***Table S6***). The associated SNPs may tag two upstream common deletions (8 and 43 kb) that delete transcription factor binding sites,^46^ although reports differ on whether the signal is driven by the shorter^27^ or the longer deletion.^31^ Thus, the top two associations are in or near genes that influence BMI and may be involved in neurite outgrowth and synaptic plasticity.

Novel associations reported here include *RBFOX1* and *LRFN5*. There are independent associations with MDD at both the 5′ and the 3′ ends of *RBFOX1* (1.7 Mb, RNA binding protein fox-1 homolog 1). This convergence makes it a strong candidate gene. Fox-1 regulates the expression of thousands of genes, many of which are expressed at synapses and enriched for autism-related genes.^47^ The Fox-1 network regulates neuronal excitability and prevents seizures.^48^ It directs splicing in the nucleus and binds to 3′ UTRs of target mRNAs in the cytoplasm.^48,49^ Of particular relevance to MDD, Fox-1 participates in the termination of the corticotropin releasing hormone response to stress by promoting alternative splicing of the PACAP receptor to its repressive form.^50^ Thus, *RBFOX1* could play a role in the chronic hypothalamic-pituitary-adrenal axis hyperactivation that has been widely reported in MDD.^51^

*LRFN5* (leucine rich repeat and fibronectin type III domain containing 5) encodes adhesion-like molecules involved in synapse formation. Common SNPs in *LRFN5* were associated with depressive symptoms in older adults in a gene-based GWA analysis.^52^ LRFN5 induces excitatory and inhibitory presynaptic differentiation in contacting axons and regulates synaptic strength.^53,54^ LRFN5 also limits T-cell response and neuroinflammation (CNS “immune privilege”) by binding to herpes virus entry mediator; a LRFN5-specific monoclonal antibody increases activation of microglia and macrophages by lipopolysaccharide and exacerbates mouse experimental acquired encephalitis;^55^ thus, reduced expression (the predicted effect of eQTLs in LD with the associated SNPs) could increase neuroinflammatory responses.

Gene-wise analyses identified 153 significant genes after controlling for multiple comparisons (***Table S7***). Many of these genes were in the extended MHC region (45 of 153) and their interpretation is complicated by high LD and gene density. In addition to the genes discussed above, other notable and significant genes outside of the MHC include multiple potentially “druggable” targets that suggest connections of the pathophysiology of MDD to neuronal calcium signaling (*CACNA1E* and *CACNA2D1*), dopaminergic neurotransmission (*DRD2*, a principal target of antipsychotics), glutamate neurotransmission (*GRIK5* and *GRM5*), and presynaptic vesicle trafficking (*PCLO*).

Finally, comparison of the MDD loci with 108 loci for schizophrenia^22^ identified six shared loci. Many SNPs in the extended MHC region are strongly associated with schizophrenia, but implication of the MHC region is novel for MDD. Another example is *TCF4* (transcription factor 4) which is strongly associated with schizophrenia but not previously with MDD. TCF4 is essential for normal brain development, and rare mutations in *TCF4* cause Pitt–Hopkins syndrome which includes autistic features.^56^ GRS calculated from the schizophrenia GWA results explained 0.8% of the variance in liability of MDD (***Figure 2c***).

## Implications for the biology of MDD using functional genomic data

Results from “-omic” studies of functional features of cells and tissues are necessary to understand the biological implications of results of GWA for complex disorders like MDD.^57^ To further elucidate the biological relevance of the MDD findings, we integrated the results with a wide range of functional genomic data. First, using enrichment analyses, we compared the MDD GWA findings to bulk tissue mRNA-seq from GTEx.^58^ Only brain samples showed significant enrichment (***Figure 3A***), and the three tissues with the most significant enrichments were all cortical. Prefrontal cortex and anterior cingulate cortex are important for higher-level executive functions and emotional regulation which are often impaired in MDD. Both regions were implicated in a large meta-analysis of brain MRI findings in adult MDD cases.^59^ Second, given the predominance of neurons in cortex, we confirmed that the MDD genetic findings connect to genes expressed in neurons but not oligodendrocytes or astrocytes (***Figure 3B***).^60^ These results confirm that MDD is a brain disorder and provide validation for the utility of our genetic results for the etiology of MDD.

**Figure 3:**
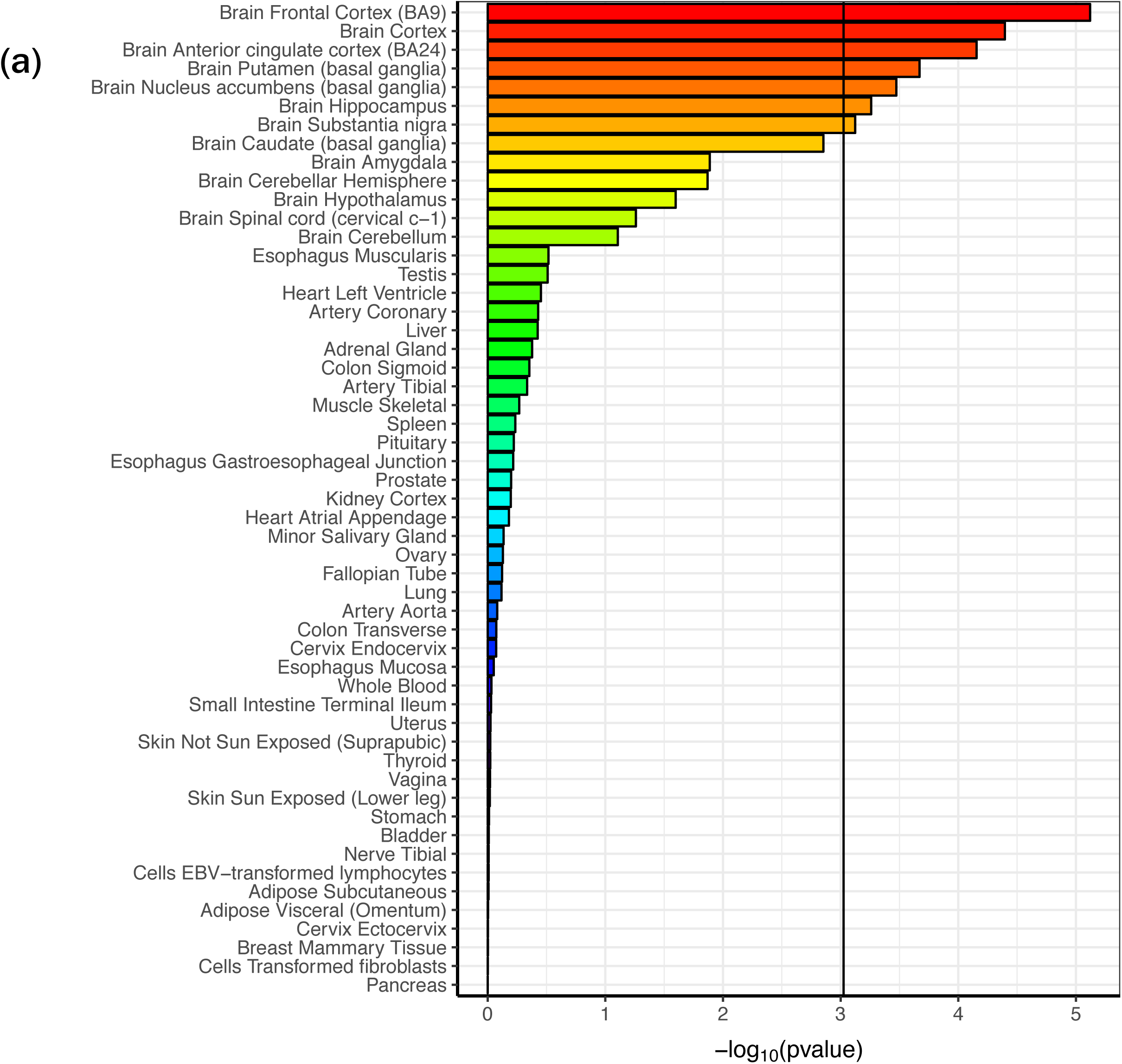
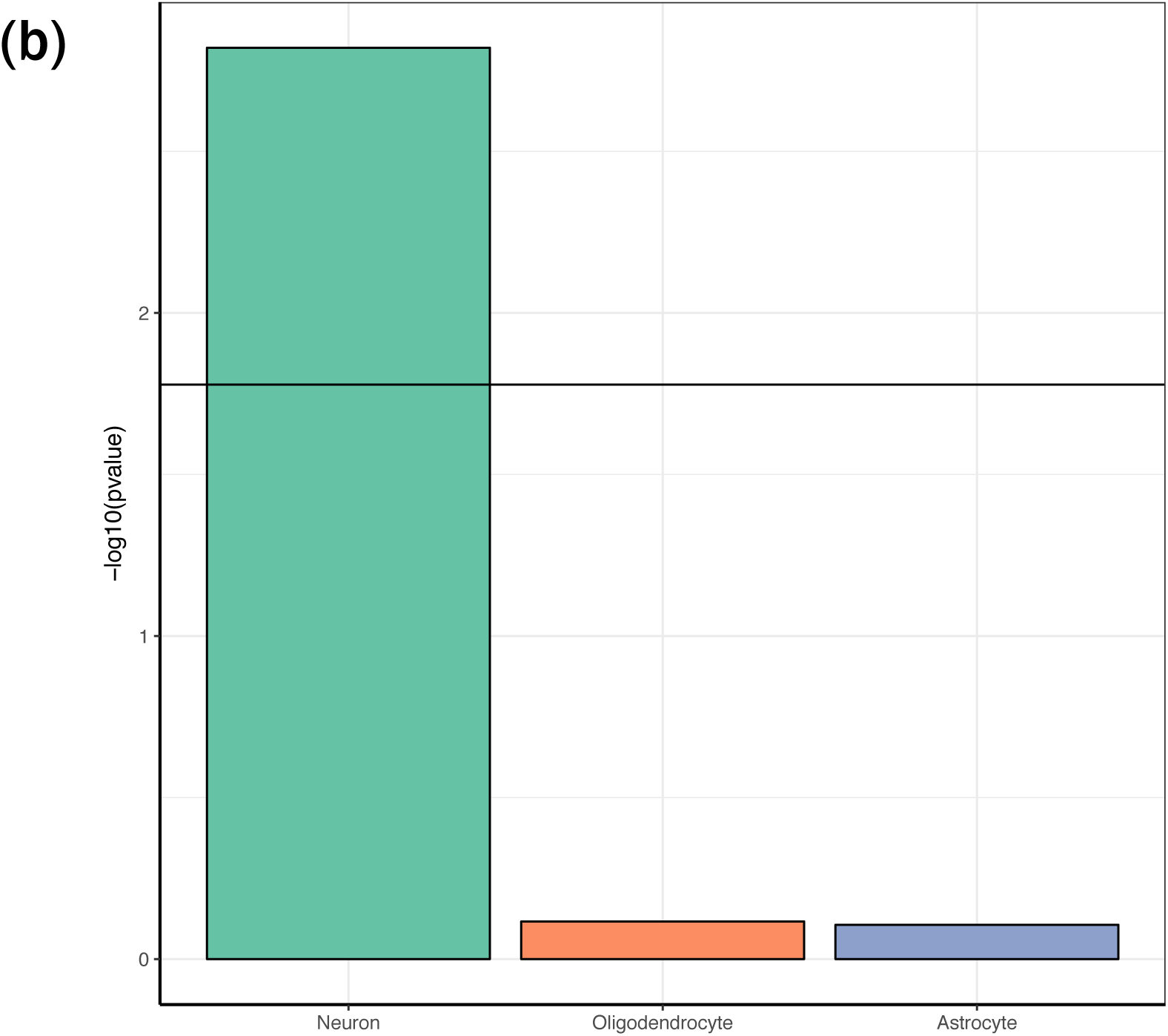
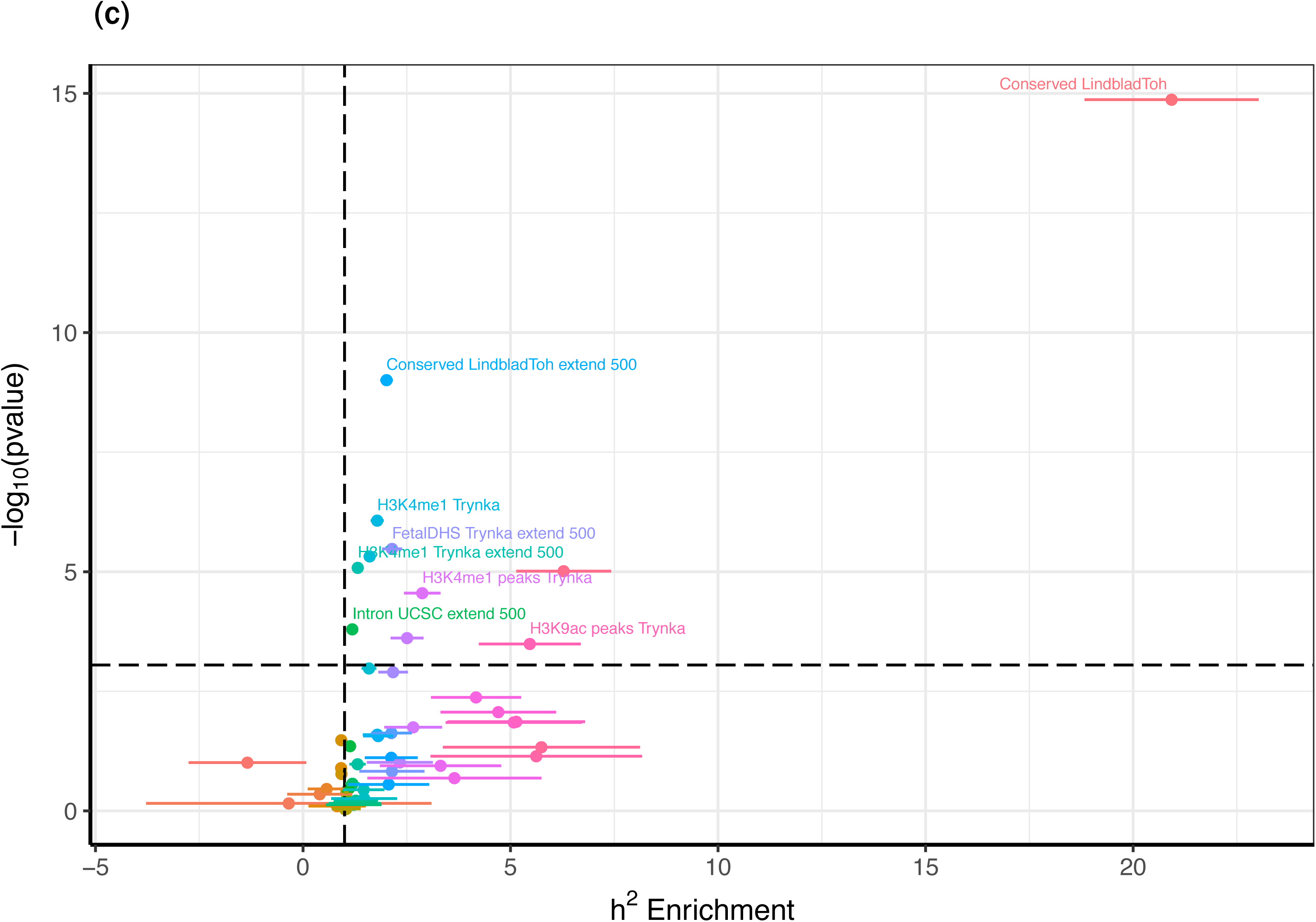
Comparisons of the MDD GWA meta-analysis. (a) MDD results and enrichment in bulk tissue mRNA-seq from GTEx. Only brain tissues showed enrichment, and the three tissues with the most significant enrichments were all cortical. (b) MDD results and enrichment in three major brain cell types. The MDD genetic findings were enriched in neurons but not oligodendrocytes or astrocytes. (c) Partitioned LDSC to evaluate enrichment of the MDD GWA findings in over 50 functional genomic annotations (**Table S8**). The major finding was the significant enrichment of MDD 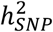 in genomic regions conserved across 29 Eutherian mammals.^62^ Other enrichments implied regulatory activity, and included open chromatin in human brain and an epigenetic mark of active enhancers (H3K4me1). Exonic regions did not show enrichment. We found no evidence that Neanderthal introgressed regions were enriched for MDD GWA findings.

Third, we used partitioned LD score regression^61^ to evaluate the enrichment of the MDD GWA findings in over 50 functional genomic annotations (***Figure 3C*** and ***Table S8***). The major finding was the significant enrichment of MDD 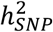 in genomic regions conserved across 29 Eutherian mammals^62^ (20.9 fold enrichment, *P*=1.4×10^−15^). This annotation was also the most enriched for schizophrenia.^61^ We could not evaluate regions conserved in primates or human “accelerated” regions as there were too few for confident evaluation.^62^ The other major enrichments implied regulatory activity, and included open chromatin in human brain and an epigenetic mark of active enhancers (H3K4me1). Notably, exonic regions did not show enrichment suggesting that, as with schizophrenia,^20^ genetic variants that change exonic sequences may not play a large role in MDD. We found no evidence that Neanderthal introgressed regions were enriched for MDD GWA findings.^63^

Fourth, we applied methods to integrate GWA SNP-MDD results with those from gene expression quantitative trait loci (eQTL) studies. SMR (summary data–based Mendelian randomization)^64^ identified 13 MDD-associated SNPs with strong evidence that they control local gene expression in one or more tissues (***Table S9*** and ***Figure S2***), including two loci not reaching GWA significance (*TMEM64* and *ZDHHC5*). A transcriptome-wide association study^65^ applied to data from the dorsolateral prefrontal cortex^66^ identified 17 genes where MDD-associated SNPs influenced gene expression (***Table S10***). These genes included *OLFM4* (discussed above).

Fifth, we added additional data types to attempt to improve understanding of individual loci. For the intergenic associations, we evaluated total-stranded RNA-seq data from human brain and found no evidence for unannotated transcripts in these regions. A particularly important data type is assessment of DNA-DNA interactions which can localize a GWA finding to a specific gene that may be nearby or hundreds of kb away.^67-69^ We integrated the MDD findings with “easy Hi-C” data from brain cortical samples (3 adult, 3 fetal, more than 1 billion reads each). These data clarified three of the associations.

The statistically independent associations in *NEGR1* (rs1432639, *P*=4.6×10^−15^) and over 200 kb away (rs12129573, *P*=4.0×10^−12^) both implicate *NEGR1* (***Figure S3a***), the former likely due to the presence of a reportedly functional copy number polymorphism (see above) and the presence of intergenic loops. The latter association has evidence of DNA looping interactions with *NEGR1*. The association in *SOX5* (rs4074723) and the two statistically independent associations in *RBFOX1* (rs8063603 and rs7198928, *P*=6.9×10^−9^ and 1.0×10^−8^) had only intragenic associations, suggesting that the genetic variation in the regions of the MDD associations act locally and can be assigned to these genes. In contrast, the association in *RERE* (rs159963 *P*=3.2×10^−8^) could not be assigned to *RERE* as it may contain super-enhancer elements given its many DNA-DNA interactions with many nearby genes (***Figure S3b***).

## Implications for the biology of MDD based on the roles of sets of genes

A parsimonious explanation for the presence of many significant associations for a complex trait like MDD is that the different associations are part of a higher order grouping of genes.^70^ These could be a biological pathway or a collection of genes with a functional connection. Multiple methods allow evaluation of the connection of MDD GWA results to sets of genes grouped by empirical or predicted function (i.e., pathway or gene set analysis).

Full pathway analyses are shown in ***Table S11***, and the 19 pathways with false discovery rate q-values < 0.05 are summarized in ***Figure 4***. The major groupings of significant pathways were: RBFOX1, RBFOX2, RBFOX3, or CELF4 regulatory networks; genes whose mRNAs are bound by FMRP; synaptic genes; genes involved in neuronal morphogenesis; genes involved in neuron projection; genes associated with schizophrenia (at *P*<10^−4^)^22^; genes involved in CNS neuron differentiation; genes encoding voltage-gated calcium channels; genes involved in cytokine and immune response; and genes known to bind to the retinoid X receptor. Several of these pathways are implicated by GWA of schizophrenia and by rare exonic variation of schizophrenia and autism,^71,72^ and immediately suggest shared biological mechanisms across these disorders.

**Figure 4:**
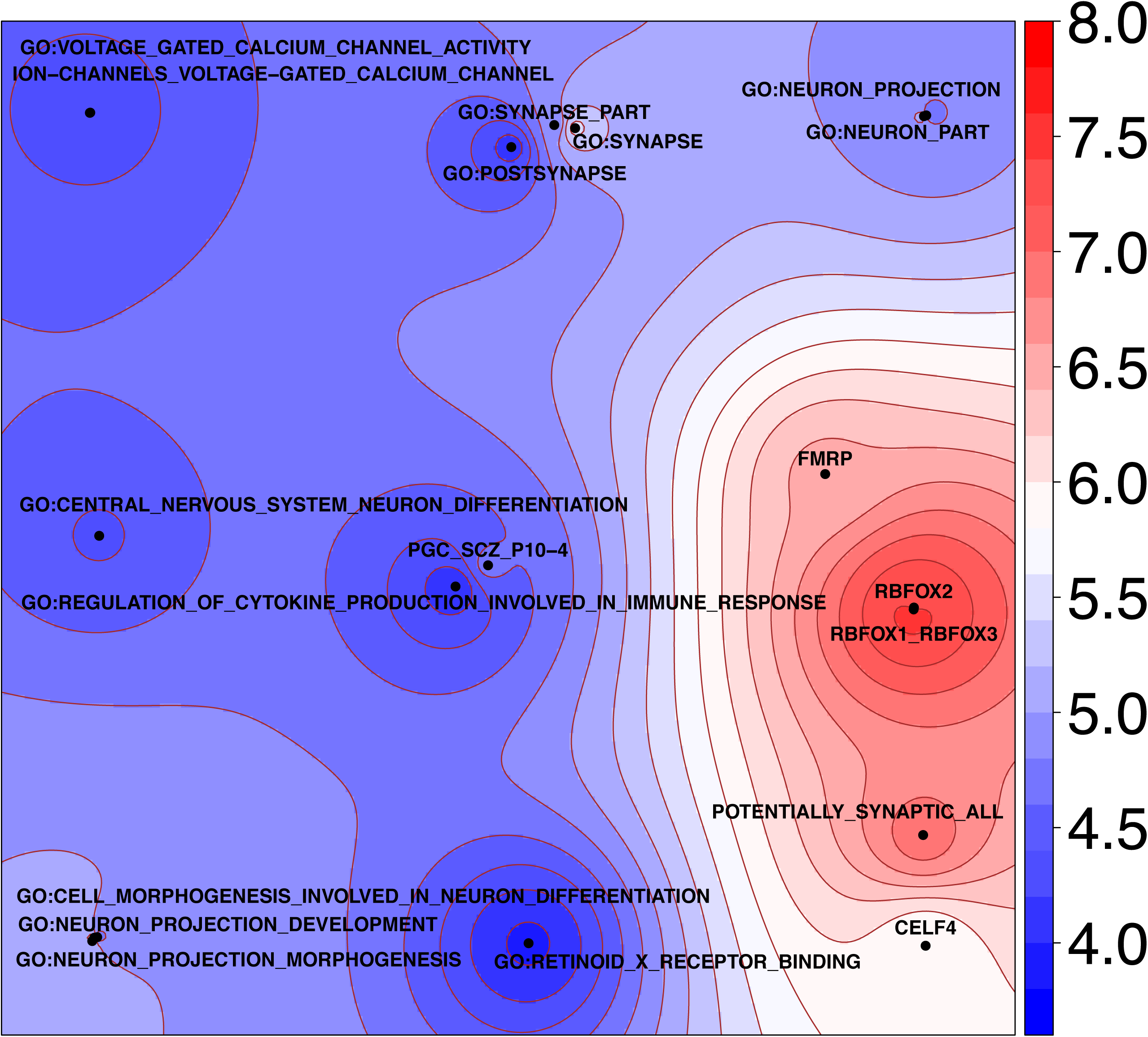
Generative topographic mapping of the 19 significant pathway results. The average position of each pathway on the map is represented by a point. The map is colored by the −log_10_(P) obtained using MAGMA. The X and Y coordinates result from a kernel generative topographic mapping algorithm (GTM) that reduces high dimensional gene sets to a two-dimensional scatterplot by accounting for gene overlap between gene sets. Each point represents a gene set. Nearby points are more similar in gene overlap than more distant points. The color surrounding each point (gene set) indicates significance per the scale on the right. The significant pathways (*Table S11*) fall into nine main clusters as described in the text.

A key issue for common variant GWA studies is their relevance for pharmacotherapy: do the results connect meaningfully to known medication targets and might they suggest new mechanisms or “druggable” targets? We conducted gene set analysis that compared the MDD GWA results to targets of antidepressant medications defined by pharmacological studies,^73^ and found that 42 sets of genes encoding proteins bound by antidepressant medications were highly enriched for smaller MDD association *P*-values than expected by chance (42 drugs, rank enrichment test *P*=8.5×10^−10^). This finding connects our MDD genomic findings to MDD therapeutics, and suggests the salience of these results for novel lead compound discovery for MDD.^74^

## Implications for a deeper understanding of the clinically-defined entity “MDD”

Past epidemiological studies associated MDD with many other diseases and traits. Due to limitations inherent to observational studies, understanding whether a phenotypic correlation is potentially causal or if it results from reverse causation or confounding is generally unclear. Genetic studies can now offer complementary strategies to assess whether a phenotypic association between MDD and a risk factor or a comorbidity is mirrored by a non-zero *r_g_* (common variant genetic correlation) and, for some of these, evaluate the potential causality of the association given that exposure to genetic risk factors begins at conception.

We used LD score regression to estimate *r_g_* of MDD with 221 psychiatric disorders, medical diseases, and human traits.^14,75^ ***Table S12*** contains the full results, and ***Table 2*** holds the *r_g_* values with false discovery rates < 0.01. First, there were very high genetic correlations for MDD with current depressive symptoms. Both correlations were close to +1 (the samples in one report overlapped partially with this MDD meta-analysis^18^ but the other did not^16^). The *r_g_* estimate in the MDD anchor samples with depressive symptoms was numerically smaller (0.80, SE 0.059) but the confidence intervals overlapped those for the full sample. Thus, the common-variant genetic architecture of lifetime MDD overlapped strongly with that of current depressive symptoms (bearing in mind that current symptoms had lower estimates of 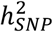 compared to the lifetime measure of MDD).

**Table 2.** Genetic correlations of MDD with other disorders, diseases, and human traits

| Trait | $r_g$ | SE | FDR | $h_{SNP}^2$ | PMID |
| --- | --- | --- | --- | --- | --- |
| Depressive symptoms, CHARGE | 0.91 | 0.123 | 3.2e-12 | 0.04 | 23290196 |
| Depressive symptoms, SSGAC | 0.98 | 0.034 | 1.3e-176 | 0.05 | 27089181 |
| ADHD (iPSYCH-PGC) | 0.42 | 0.033 | 6.1e-36 | 0.24 | submitted |
| Anorexia nervosa | 0.13 | 0.028 | 7.1e-5 | 0.55 | 24514567 |
| Anxiety disorders | 0.80 | 0.140 | 2.0e-7 | 0.06 | 26857599 |
| Autism spectrum disorders (iPSYCH-PGC) | 0.44 | 0.039 | 8.4e-28 | 0.20 | submitted |
| Bipolar disorder | 0.32 | 0.034 | 3.3e-19 | 0.43 | 21926972 |
| Schizophrenia | 0.34 | 0.025 | 7.7e-40 | 0.46 | 25056061 |
| Smoking, ever vs never | 0.29 | 0.038 | 7.0e-13 | 0.08 | 20418890 |
| Daytime sleepiness ‡ | 0.19 | 0.048 | 5.7e-4 | 0.05 | 0 |
| Insomnia ‡ | 0.38 | 0.038 | 4.0e-22 | 0.13 | 0 |
| Tiredness | 0.67 | 0.037 | 6.2E-72 | 0.07 | 28194004 |
| Subjective well-being | -0.65 | 0.035 | 7.5E-76 | 0.03 | 27089181 |
| Neuroticism | 0.70 | 0.031 | 2.5E-107 | 0.09 | 27089181 |
| College completion | -0.17 | 0.034 | 6.7E-6 | 0.08 | 23722424 |
| Years of education | -0.13 | 0.021 | 1.6E-8 | 0.13 | 27225129 |
| Body fat | 0.15 | 0.038 | 6.5e-4 | 0.11 | 26833246 |
| Body mass index | 0.09 | 0.026 | 3.6e-3 | 0.19 | 20935630 |
| Obesity class 1 | 0.11 | 0.029 | 1.6e-3 | 0.22 | 23563607 |
| Obesity class 2 | 0.12 | 0.033 | 3.0e-3 | 0.18 | 23563607 |
| Obesity class 3 | 0.20 | 0.053 | 1.6e-3 | 0.12 | 23563607 |
| Overweight | 0.13 | 0.030 | 1.4e-4 | 0.11 | 23563607 |
| Waist circumference | 0.11 | 0.024 | 8.2e-5 | 0.12 | 25673412 |
| Waist-to-hip ratio | 0.12 | 0.030 | 2.9e-4 | 0.11 | 25673412 |
| Triglycerides | 0.14 | 0.028 | 1.0e-5 | 0.17 | 20686565 |
| Age at menarche | -0.14 | 0.023 | 6.3E-8 | 0.20 | 25231870 |
| Age of first birth | -0.29 | 0.029 | 6.1E-22 | 0.06 | 27798627 |
| Fathers age at death | -0.28 | 0.058 | 3.0E-5 | 0.04 | 27015805 |
| Number of children ever born | 0.13 | 0.036 | 2.4E-3 | 0.03 | 27798627 |
| Coronary artery disease | 0.12 | 0.027 | 8.2e-5 | 0.08 | 26343387 |
| Squamous cell lung cancer | 0.26 | 0.075 | 3.6e-3 | 0.04 | 27488534 |

‡ Self-reported daytime sleepiness and insomnia from UK Biobank excluding subjects with MDD, other psychiatric disorders (bipolar disorder, schizophrenia, autism, intellectual disability), shift workers, and those taking hypnotics.

Second, MDD had significant positive genetic correlations with every psychiatric disorder assessed as well as with smoking initiation. This is the most comprehensive and best-powered evaluation of the relation of MDD with other psychiatric disorders yet published, and these results indicate that the common genetic variants that predispose to MDD overlap substantially with those for adult and childhood onset psychiatric disorders.

Third, MDD had positive genetic correlations with multiple measures of sleep quality (daytime sleepiness, insomnia, and tiredness). The first two of these correlations were based on a specific analysis of UK Biobank data (i.e., removing people with MDD, other major psychiatric disorders, shift workers, and those taking hypnotics). This pattern of correlations combined with the critical importance of sleep and fatigue in MDD (these are two commonly accepted criteria for MDD) suggests a close and potentially profound mechanistic relation. MDD also had a strong genetic correlation with neuroticism (a personality dimension assessing the degree of emotional instability); this is consistent with the literature showing a close interconnection of MDD and this personality trait. The strong negative *r_g_* with subjective well-being underscores the capacity of MDD to impact human health.

Finally, MDD had negative correlations with two proxy measures of intelligence, positive correlations with multiple measures of adiposity, relationship to female reproductive behavior (decreased age at menarche, age at first birth, and increased number of children), and positive correlations with coronary artery disease and lung cancer.

We used Mendelian randomization (MR) to investigate the relationships between genetically correlated traits. We conducted bi-directional MR analysis for four traits: years of education (EDY, a proxy for general intelligence)^76^, body mass index (BMI)^27^, coronary artery disease (CAD)^77^, and schizophrenia^22^. These traits were selected because all of the following were true: phenotypically associated with MDD, significant *r_g_* with MDD with an unclear direction of causality, and >30 independent genome-wide significant associations from large GWA.

We report GSMR (generalized summary statistic-based MR) results but obtained qualitatively similar results with other MR methods (***Table S13*** and *Figures S4A-D*). MR analyses provided evidence for a 1.15-fold increase in MDD per standard deviation of BMI (*P*_GSMR_=2.7×10^−7^) and a 0.89-fold decrease in MDD per standard deviation of EDY (*P*_GSMR_=8.8×10^−7^). There was no evidence of reverse causality of MDD for BMI (*P*_GSMR_=0.81) or EDY (*P*_GSMR_=0.28). For BMI there was some evidence of pleiotropy, as eight SNPs were excluded by the HEIDI-outlier test including SNPs near *OLFM4* and *NEGR1* (if these were included, the estimate of increased risk for MDD was greater). Thus, these results are consistent with EDY and BMI as causal risk factors or correlated with causal risk factors for MDD. For CAD, the MR analyses were not significant when considering MDD as an outcome (*P*_GSMR_=0.39) or as an exposure (*P*_GSMR_=0.13). We interpret the *r_g_* of 0.12 between CAD and MDD to reflect a genome-wide correlation in the sign of effect sizes but no correlation in the effect size magnitudes: this is consistent with “type I pleiotropy”^78^, that there are multiple molecular functions of these genetic variants (which may be tissue-specific in brain and heart). However, because the MR regression coefficient for MDD instruments has relatively high standard error, this analysis should be revisited when more MDD genome-wide significant SNP instruments become available from future MDD GWA studies.

We used MR to investigate the relationship between MDD and schizophrenia. Although MDD had positive *r_g_* with many psychiatric disorders, only schizophrenia has sufficient associations for MR analyses. We found significant bi-directional correlations in SNP effect sizes for schizophrenia loci in MDD (*P*_GSMR_=7.7×10^−46^) and for MDD loci in schizophrenia (*P*_GSMR_=6.3×10^−15^). We interpret the MDD-schizophrenia *r_g_* of 0.34 as reflecting type II pleiotropy^78^ (i.e., consistent with shared biological pathways being causal for both disorders).

## Empirically, what is MDD?

The nature of severe depression has been discussed for millennia.^79^ This GWA meta-analysis is among the largest ever conducted for a psychiatric disorder, and provides a body of results that help refine and define the fundamental basis of MDD.

First, MDD is a brain disorder. Although this is not unexpected, some past models of MDD have had little or no place for heredity or biology. Our results indicate that genetics and biology are definite pieces in the puzzle of MDD. The genetic results best match gene expression patterns in prefrontal and anterior cingulate cortex, anatomical regions that show differences between MDD cases and controls. The genetic findings implicated neurons (not microglia or astrocytes), and we anticipate more detailed cellular localization when sufficient single-cell and single-nuclei RNA-seq datasets become available.^80^

Second, the genetic associations for MDD (as with schizophrenia)^61^ tend to occur in genomic regions conserved across a range of placental mammals. Conservation suggests important functional roles. Given that this analysis did not implicate exons or coding regions, MDD may not be characterized by common changes in the amino acid content of proteins.

Third, the results also implicated developmental gene regulatory processes. For instance, the genetic findings pointed at *RBFOX1* (the presence of two independent genetic associations in *RBFOX1* strongly suggests that it is the MDD-relevant gene). Gene set analyses implicated genes containing binding sites to the protein product of *RBFOX1* in MDD, and this gene set is also significantly enriched for rare exonic variation in autism and schizophrenia.^71,72^ These analyses highlight the potential importance of splicing to generate alternative isoforms; risk for MDD may be mediated not by changes in isolated amino acids but rather by changes in the proportions of isoforms coming from a gene, given that isoforms often have markedly different biological functions.^81,82^ These convergent results provide a tantalizing suggestion of a biological mechanism common to multiple severe psychiatric disorders.

Fourth, in the most extensive analysis of the genetic “connections” of MDD with a wide range of disorders, diseases, and human traits, we found significant positive genetic correlations with measures of body mass and negative genetic correlations with years of education. MR analyses suggested the *potential* causality of both correlations, and our results certainly provide hypotheses for more detailed prospective studies. However, further clarity requires larger and more informative GWA studies for a wider range of related traits (e.g., with >30 significant associations per trait). We strongly caution against interpretations of these results that go beyond the analyses undertaken (e.g., these results do not provide evidence that weight loss would have an antidepressant effect). The currently available data do not provide further insight about the fundamental driver or drivers of causality. The underlying mechanisms are likely more complex as it is difficult to envision how genetic variation in educational attainment or body mass alters risk for MDD without invoking an additional mechanistic component. For example, genetic variation underlying general intelligence might directly alter the development and function of discrete brain regions that alters intelligence and which also predisposes to worse mood regulation. Alternatively, genetic variation underlying general intelligence might lead to poorer development of cognitive strategies to handle adversity which increases risk for MDD. An additional possibility is that there are sets of correlated traits–e.g., personality, intelligence, sleep patterns, appetitive regulation, or propensity to exercise–and that these act in varying combinations in different people. Our results are inconsistent with a causal relation between MDD and subsequent changes in body mass or education years. If such associations are observed in epidemiological or clinical samples, then it is likely not MDD but something correlated with MDD that drives the association.

Fifth, we found significant positive correlations of MDD with all psychiatric disorders that we evaluated, including disorders prominent in childhood. This pattern of results indicates that the current classification scheme for major psychiatric disorders does not align well with the underlying genetic basis of these disorders. The MR results for MDD and schizophrenia indicated a shared biological basis.

The dominant psychiatric nosological systems were principally designed for clinical utility, and are based on data that emerge during human interactions (i.e., observable signs and reported symptoms) and not objective measurements of pathophysiology. MDD is frequently comorbid with other psychiatric disorders, and the phenotypic comorbidity has an underlying structure that reflects shared origins (as inferred from factor analyses and twin studies).^83-86^ Our genetic results add to this knowledge: MDD is not a discrete entity at any level of analysis. Rather, our data strongly suggest the existence of biological processes common to MDD and schizophrenia. It would be unsurprising if future work implicated bipolar disorder, anxiety disorders, and other psychiatric disorders as well.

Finally, as expected, we found that MDD had modest 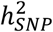 (8.9%) since MDD is a complex malady with both genetic and environmental determinants. We found that MDD has a very high genetic correlation with proxy measures that can be briefly assessed. Lifetime major depressive disorder requires a constellation of signs and symptoms whose reliable scoring requires an extended interview with a trained clinician. However, the common variant genetic architecture of lifetime major depressive <u>disorder</u> in these seven cohorts (containing many subjects medically treated for MDD) has strong overlap with that of current depressive <u>symptoms</u> in general community samples. Similar relations of clinically-defined ADHD or autism with quantitative genetic variation in the population have been reported.^87,88^ The MDD “disorder versus symptom” relationship has been debated extensively,^89^ but our data indicate that the common variant genetic overlap is very high. This finding has two important implications.

One implication is for future genetic studies of MDD. In a first phase, it should be possible to elucidate the bulk of the common variant genetic architecture of MDD using a cost-effective shortcut – large studies of genotyped individuals who complete brief lifetime MDD screening (a sample size approaching 1 million MDD cases may be achievable by 2020). In a second phase, with a relatively complete understanding of the genetic basis of MDD, one could then evaluate smaller samples of carefully phenotyped individuals with MDD to understand the clinical importance of the genetic results. These data could allow more precise delineation of the clinical heterogeneity of MDD (e.g., our demonstration that individuals with more severe or recurrent MDD have inherited a higher genetic loading for MDD than single-episode MDD). Subsequent empirical studies may show that it is possible to stratify MDD cases at first presentation to identify individuals at high risk for recurrence, poor outcome, poor treatment response, or who might subsequently develop a psychiatric disorder requiring alternative pharmacotherapy (e.g., schizophrenia or bipolar disorder). This could form a cornerstone of precision medicine in psychiatry.

The second implication is that people with MDD differ only by degree from those who have not experienced MDD. All humans carry lesser or greater numbers of genetic risk factors for MDD. Genetic risk for MDD is continuous and normally distributed with no clear point of demarcation. Non-genetic factors play important protective and pre-disposing roles (e.g., life events, exposure to chronic fear, substance abuse, and a wide range of life experiences and choices). The relation of blood pressure to essential hypertension is a reasonable analogy. All humans inherit different numbers of genetic variants that influence long-term patterns of blood pressure with environmental exposures and life choices also playing roles. The medical “disorder” of hypertension is characterized by blood pressure chronically over a numerical threshold above which the risks for multiple preventable diseases climb. MDD is not a “disease” (i.e., a distinct entity delineable using an objective measure of pathophysiology) but indeed a disorder, a human-defined but definable syndrome that carries increased risk of adverse outcomes. The adverse outcomes of hypertension are diseases (e.g., stroke or myocardial infarction). The adverse outcomes of MDD include elevation in risk for a few diseases, but the major impacts of MDD are death by suicide and disability.

In summary, this GWA meta-analysis of 130,664 MDD cases and 330,470 controls identified 44 loci. An extensive set of companion analyses provide insights into the nature of MDD as well as its neurobiology, therapeutic relevance, and genetic and biological interconnections to other psychiatric disorders. Comprehensive elucidation of these features is the primary goal of our genetic studies of MDD.

## Online Methods

### Anchor cohort

Our analysis was anchored in a GWA mega-analysis of 29 samples of European-ancestry (16,823 MDD cases and 25,632 controls). ***Table S1*** summarizes the source and inclusion/exclusion criteria for cases and controls for each sample. All samples in the initial PGC MDD papers were included.^13,15,90^ All anchor samples passed a structured methodological review by MDD assessment experts (DF Levinson and KS Kendler). Cases were required to meet international consensus criteria (DSM-IV, ICD-9, or ICD-10)^91-93^ for a lifetime diagnosis of MDD established using structured diagnostic instruments from assessments by trained interviewers, clinician-administered checklists, or medical record review. All cases met standard criteria for MDD, were directly interviewed (28/29 samples) or had medical record review by an expert diagnostician (1/29 samples), and most were ascertained from clinical sources (19/29 samples). Controls in most samples were screened for the absence of lifetime MDD (22/29 samples), and randomly selected from the population. We considered this the “anchor” cohort given use of standard methods of establishing the presence or absence of MDD.

The most direct and important way to evaluate the comparability of the samples comprising the anchor cohort is using SNP genotype data.^14,94^ The sample sizes were too small to evaluate the common variant genetic correlations (*r_g_*) between all pairs of anchor cohort samples (>3,000 subjects per sample are recommended). As an alternative, we used “leave one out” genetic risk scores (GRS, described below). We repeated this procedure by leaving out each of the anchor cohort samples so that we could evaluate the similarity of the common-variant genetic architectures of each sample to the rest of the anchor cohort. ***Figure S1A*** shows that all samples in the anchor cohort (except one) yielded significant differences in case-control distributions of GRS.

### Expanded cohorts

We critically evaluated an “expanded” set of six independent, European-ancestry cohorts (113,841 MDD cases and 304,838 controls). ***Table S2*** summarizes the source and inclusion/exclusion criteria for cases and controls for each cohort. These cohorts used a range of methods for assessing MDD: Generation Scotland employed direct interviews; iPSYCH (Denmark) used national treatment registers; deCODE (Iceland) used national treatment registers and direct interviews; GERA used Kaiser-Permanente treatment records (CA, US); UK Biobank combined self-reported MDD symptoms and/or treatment for MDD by a medical professional; and 23andMe used self-report of treatment for MDD by a medical professional. All controls were screened for the absence of MDD.

### Cohort comparability

***Table S3*** summarizes the numbers of cases and controls in the anchor cohort and the six expanded cohorts. The most direct and important way to evaluate the comparability of these cohorts for a GWA meta-analysis is using SNP genotype data.^14,94^ We used LD score regression (described below) to estimate 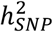 for each cohort, and *r_g_* for all pairwise combinations of the cohorts.

We compared the seven anchor and expanded cohorts. First, there was no indication of important sample overlap as the LDSC regression intercept between pairs of cohorts ranged from −0.01 to +0.01. Second, ***Table S4*** shows 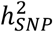 on the liability scale for each cohort. The 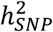 estimates range from 0.09 to 0.23 (for lifetime risk (=0.15) but the confidence intervals largely overlap. Third, ***Table S4*** also shows the *r_g_* values for all pairs of anchor and expanded cohorts. The median *r_g_* was 0.80 (interquartile range 0.67–0.96), and the upper 95% confidence interval on *r_g_* included 0.75 for all pairwise comparisons. These results indicate that the common variant genetic architecture of the anchor and expanded cohorts overlap strongly, and provide critical support for the full meta-analysis of all cohorts.

### Genotyping and quality control

Genotyping procedures can be found in the primary reports for each cohort (***Tables S1-S2***). Individual genotype data for all anchor cohorts, GERA, and iPSYCH were processed using the PGC “ricopili” pipeline (URLs) for standardized quality control, imputation, and analysis.^22^ The expanded cohorts from deCODE, Generation Scotland, UK Biobank, and 23andMe were processed by the collaborating research teams using comparable procedures. SNPs and insertion-deletion polymorphisms were imputed using the 1000 Genomes Project multi-ancestry reference panel (URLs).^95^

Quality control and imputation on the 29 PGC MDD anchor cohorts was performed according to standards from the PGC (***Table S3***). The default parameters for retaining SNPs and subjects were: SNP missingness < 0.05 (before sample removal); subject missingness < 0.02; autosomal heterozygosity deviation (|*F_het_*|<0.2); SNP missingness < 0.02 (after sample removal); difference in SNP missingness between cases and controls < 0.02; and SNP Hardy-Weinberg equilibrium (*P* > 10^−6^ in controls or *P* > 10^−10^ in cases). These default parameters sufficiently controlled λ and false positive findings for 16 cohorts (boma, rage, shp0, shpt, edi2, gens, col3, mmi2, qi3c, qi6c, qio2, rai2, rau2, twg2, grdg, grnd). Two cohorts (gep3 and nes2) needed stricter SNP filtering and 11 cohorts needed additional ancestral matching (rot4, stm2, rde4) or ancestral outlier exclusion (rad2, i2b3, gsk1, pfm2, jjp2, cof3, roc3, mmo4). An additional cohort of inpatient MDD cases from Münster, Germany was processed through the same pipeline.

Genotype imputation was performed using the pre-phasing/imputation stepwise approach implemented in IMPUTE2 / SHAPEIT (chunk size of 3 Mb and default parameters). The imputation reference set consisted of 2,186 phased haplotypes from the 1000 Genomes Project dataset (August 2012, 30,069,288 variants, release “v3.macGT1”). After imputation, we identified SNPs with very high imputation quality (INFO >0.8) and low missingness (<1%) for building the principal components to be used as covariates in final association analysis. After linkage disequilibrium pruning (r^2^ > 0.02) and frequency filtering (MAF > 0.05), there were 23,807 overlapping autosomal SNPs in the data set. This SNP set was used for robust relatedness testing and population structure analysis. Relatedness testing identified pairs of subjects with 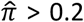, and one member of each pair was removed at random after preferentially retaining cases over controls. Principal component estimation used the same collection of autosomal SNPs.

Identification of identical samples is easily accomplished given direct access to individual genotypes.^13^ Two concerns are the use of the same control samples in multiple studies (e.g., GAIN or WTCCC controls)^96,97^ and inclusion of closely related individuals. For cohorts where the PGC central analysis team had access to individual genotypes (all anchor cohorts and GERA), we used SNPs directly genotyped on all platforms to compute empirical relatedness, and excluded one of each duplicated or relative pair (defined as 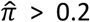). Within all other cohorts (deCODE, Generation Scotland, iPSYCH, UK Biobank, 23andMe, and CONVERGE), identical and relative pairs were identified and resolved using similar procedures. Identical samples between the anchor cohorts, iPSYCH, UK Biobank, and Generation Scotland were identified using genotype-based checksums (URLs),^98^ and an individual on the collaborator’s side was excluded. Checksums were not available for the deCODE and 23andMe cohorts. Related pairs are not detectable by the checksum method but we did not find evidence of important overlap using LD score regression (the intercept between pairs of cohorts ranged from −0.01 to +0.01 with no evidence of important sample overlap).

### Statistical analysis

In each cohort, logistic regression association tests were conducted for imputed marker dosages with principal components covariates to control for population stratification. Ancestry was evaluated using principal components analysis applied to directly genotyped SNPs.^99^ In the anchor cohorts and GERA, we determined that all individuals in the final analyses were of European ancestry. European ancestry was confirmed in the other expanded cohorts by the collaborating research teams using similar procedures. We tested 20 principal components for association with MDD and included five principal components covariates for the anchor cohorts and GERA (all other cohorts adopted similar strategies). There was no evidence of stratification artifacts or uncontrolled test statistic inflation in the results from each anchor and extended cohort (e.g., λ_GC_ was 0.995-1.043 in the anchor cohorts). The results were combined across samples using an inverse-weighted fixed effects model.^100^ Reported SNPs have imputation marker INFO score ≥ 0.6 and allele frequencies ≥0.01 and ≤0.99, and effective sample size equivalent to > 100,000 cases. For all cohorts, X-chromosome association results were conducted separately by sex, and then meta-analysed across sexes.^22^ For two cohorts (GenScot and UKBB), we first conducted association analysis for genotyped SNPs by sex, then imputed association results using LD from the 1000 Genomes reference sample.^101^

### Defining loci

GWA findings implicate genomic regions containing multiple significant SNPs (“loci”). There were almost 600 SNPs with *P* < 5×10^−8^ in this analysis. These are not independent associations but result from LD between SNPs. We collapsed the significant SNPs to 44 loci via the following steps.

- All SNPs were high-quality (imputation INFO score ≥ 0.6 and allele frequencies ≥0.01 and ≤0.99).
- We used “clumping” to convert MDD-associated SNPs to associated regions. We identified an index SNP with the smallest *P*-value in a genomic window and other SNPs in high LD with the index SNP using PLINK (–clump-p1 1e-4 –clump-p2 1e-4 –clump-r2 0.1 –clump-kb 3000). This retained SNPs with association *P* < 0.0001 and r^2^ < 0.1 within 3 Mb windows. Only one SNP was retained from the extended MHC region due to its exceptional LD.
- We used bedtools (URLs) to combine partially or wholly overlapping clumps within 50 kb.
- We reviewed all regional plots, and removed two singleton associations (i.e., only one SNP exceeding genome-wide significance).
- We reviewed forest plots, and confirmed that association signals arose from the majority of the cohorts.
- We conducted conditional analyses. To identify independent associations within a 10 Mb region, we re-evaluated all SNPs in a region conditioning on the most significantly associated SNP using summary statistics^25^ (superimposing the LD structure from the Atherosclerosis Risk in Communities Study sample).

### Genetic risk score (GRS) analyses

To demonstrate the validity of our GWAS results, we conducted a series of GRS prediction analyses. The MDD GWA summary statistics identified associated SNP alleles and effect size which were used to calculate GRS for each individual in a target sample (i.e., the sum of the count of risk alleles weighted by the natural log of the odds ratio of the risk allele). In some analyses the target sample had been included as one of the 29 samples in the MDD anchor cohort; here, the discovery samples were meta-analyzed excluding this cohort. As in the PGC schizophrenia report,^22^ we excluded uncommon SNPs (MAF < 0.1), low-quality variants (imputation INFO < 0.9), indels, and SNPs in the extended MHC region (chr6:25-34 Mb). We then LD pruned and “clumped” the data, discarding variants within 500 kb of, and in LD r^2^ > 0.1 with the most associated SNP in the region. We generated GRS for individuals in target subgroups for a range of *P*-value thresholds (*P_T_*: 5×10^−8^, 1×10^−6^, 1×10^−4^, 0.001, 0.01, 0.05, 0.1, 0.2, 0.5, 1.0).

For each GRS analysis, five ways of evaluating the regression of phenotype on GRS are reported (***Table S5***). The significance of the case-control score difference from logistic regression including ancestry PCs and a study indicator (if more than one target dataset was analyzed) as covariates. 2) The proportion of variance explained (Nagelkerke’s R^2^) computed by comparison of a full model (covariates + GRS) to a reduced model (covariates only). It should be noted that these estimates of R^2^ reflect the proportion of cases in the case-control studies where this proportion may not reflect the underlying risk of in the population. 3) The proportion of variance on the liability scale explained by the GRS R^2^ was calculated from the difference between full and reduced linear models and was then converted to the liability scale of the population assuming lifetime MDD risk of 15%. These estimates should be comparable across target sample cohorts, whatever the proportion of cases in the sample. 4) Area under the receiver operator characteristic curve (AUC; R library pROC) was estimated in a model with no covariates^22^ where AUC can be interpreted as the probability of a case being ranked higher than a control. 5) Odds ratio for 10 GRS decile groups (these estimates also depend on both risk of MDD in the population and proportion of cases in the sample). We evaluated the impact of increasing sample size of the discovery sample GWA (***Figure 2a***) and also using the schizophrenia GWA study^22^ as the discovery sample. We also undertook GRS analysis for a target sample of MDD cases and controls not included in the meta-analysis (a clinical inpatient cohort of MDD cases and screened controls collected in Münster, Germany).

We conducted GRS analyses based on prior hypotheses from epidemiology of MDD using clinical measures available in some cohorts (if needed, the target sample was removed from the discovery GWA). We used GRS constructed from *P_T_*=0.05, selected as a threshold that gave high variance explained across cohorts (***Figure S1a***). First, we used GRS analyses to test for higher mean GRS in cases with younger age at onset (AAO) of MDD compared to those with older AAO in the anchor cohort samples. To combine analyses across samples, we used within-sample standardized GRS residuals after correcting for ancestry principal components. Heterogeneity in AAO in the anchor samples has been noted,^102^ which may reflect study specific definitions of AAO (e.g., age at first symptoms, first visit to general practitioner, or first diagnosis). Following Power et al.,^102^ we divided AAO into octiles within each cohort and combined the first three octiles into the early AAO group and the last three octiles into the late AAO group. Second, we tested for higher mean GRS for cases in anchor cohort samples with clinically severe MDD (endorsing ≥8 of 9 DSM MDD criteria) compared to those with “moderate” MDD (endorsing 5-7 of MDD criteria) following Verduijn et al.^103^ Sample sizes are given in ***Table S3***. Third, using iPSYCH as the target sample, we tested for higher mean GRS in recurrent MDD cases (ICD-10 F33, N=5,574) compared to those with single episode MDD cases (ICD-10 F32, N=12,968) in analyses that included ancestry principal components and genotyping batch as covariates. Finally, following Verduijn et al.^103^ using the NESDA sample (PGC label “nes1”, an ongoing longitudinal study of depressive and anxiety disorders) as the target sample, we constructed clinical staging phenotypes in which cases were allocated to one of three stages: Stage 2 (n = 388) first episode MDD; stage 3 (n = 562) recurrent/relapse episode MDD; stage 4 (n = 705) persistent/unremitting chronic MDD, with an episode lasting longer than 2 years before baseline interview and/or ≥ 80% of the follow-up time with depressive symptoms. We tested for higher mean GRS in stage IV cases compared to stage II MDD cases.

<u>Linkage disequilibrium (LD) score regression</u>^14,94^ was used to estimate 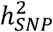 from GWA summary statistics. Estimates of 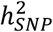 on the liability scale depend on the assumed lifetime prevalence of MDD in the population (*K*), and we assumed *K*=0.15 but also evaluated *K*=0.10 to explore sensitivity (***Table S4***). LD score regression bivariate genetic correlations attributable to genome-wide SNPs (*r_g_*) were estimated across MDD cohorts and between the full MDD cohort and other traits and disorders.

LD score regression was also used to partition 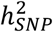 by genomic features.^61,94^ We tested for enrichment of 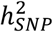 based on genomic annotations partitioning 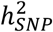 proportional to bp length represented by each annotation. We used the “baseline model” which consists of 53 functional categories. The categories are fully described elsewhere,^61^ and included conserved regions^62^, USCC gene models (exons, introns, promoters, UTRs), and functional genomic annotations constructed using data from ENCODE^104^ and the Roadmap Epigenomics Consortium.^105^ We complemented these annotations by adding introgressed regions from the Neanderthal genome in European populations^106^ and open chromatin regions from the brain dorsolateral prefrontal cortex. The open chromatin regions were obtained from an ATAC-seq experiment performed in 288 samples (N=135 controls, N=137 schizophrenia, N=10 bipolar, and N=6 affective disorder).^107^ Peaks called with MACS^108^ (1% FDR) were retained if their coordinates overlapped in at least two samples. The peaks were re-centered and set to a fixed width of 300bp using the diffbind R package.^109^ To prevent upward bias in heritability enrichment estimation, we added two categories created by expanding both the Neanderthal introgressed regions and open chromatin regions by 250bp on each side.

We used LD score regression to estimate *r_g_* between MDD and a range of other disorders, diseases, and human traits.^14^ The intent of these comparisons was to evaluate the extent of shared common variant genetic architectures in order to suggest hypotheses about the fundamental genetic basis of MDD (given its extensive comorbidity with psychiatric and medical conditions and its association with anthropometric and other risk factors). Subject overlap of itself does not bias *r_g_*.^14^ These *r_g_* are mostly based on studies of independent subjects and the estimates should be unbiased by confounding of genetic and non-genetic effects (except if there is genotype by environment correlation). When GWA studies include overlapping samples, *r_g_* remains unbiased but the intercept of the LDSC regression is an estimate of the correlation between association statistics attributable to sample overlap. These calculations were done using the internal PGC GWA library and with LD-Hub (URLs).^75^

### Relation of MDD GWA findings to tissue and cellular gene expression

We used partitioned LD score regression to evaluate which somatic tissues were enriched for MDD heritability.^110^ Gene expression data generated using mRNA-seq from multiple human tissues were obtained from GTEx v6p (URLs). Genes for which <4 samples had at least one read count per million were discarded, and samples with <100 genes with at least one read count per million were excluded. The data were normalized, and a t-statistic was obtained for each tissue by comparing the expression in each tissue with the expression of all other tissues with the exception of tissues related to the tissue of interest (e.g., brain cortex vs all other tissues excluding other brain samples), using sex and age as covariates. A t-statistic was also obtained for each tissue among its related tissue (ex: cortex vs all other brain tissues) to test which brain region was the most associated with MDD, also using sex and age as covariates. The top 10% of the genes with the most extreme t-statistic were defined as tissue specific. The coordinates for these genes were extended by a 100kb window and tested using LD score regression. Significance was obtained from the coefficient z-score, which corrects for all other categories in the baseline model.

Lists of genes specifically expressed in neurons, astrocytes, and oligodendrocytes were obtained from Cahoy et al.^60^ As these experiment were done in mice, genes were mapped to human orthologous genes using ENSEMBL. The coordinates for these genes were extended by a 100kb window and tested using LD score regression as for the GTEx tissue specific genes.

We conducted eQTL look-ups of the most associated SNPs in each region and report (***Table S6***) GWA SNPs in LD (r^2^ > 0.8) with the top eQTLs in the following data sets: eQTLGen Consortium (lllumina arrays in whole blood N=14,115, in preparation), BIOS (RNA-seq in whole blood (N=2,116),^111^ NESDA/NTR (Affymetrix arrays in whole blood, N=4,896),^112^ GEUVADIS (RNA-seq in LCL (N=465),^113^ Rosmap (RNA seq in cortex, N=494, submitted), GTEx (RNA-seq in 44 tissues, N>70),^58^ and Common Mind Consortium (CMC, prefrontal cortex, Sage Synapse accession syn5650509, N=467).^66^

We used summary-data-based Mendelian randomization (SMR)^64^ to identify loci with strong evidence of causality via gene expression (***Table S9***). SMR analysis is limited to significant cis SNP-expression (FDR < 0.05) and SNPs with MAF > 0.01 at a Bonferroni-corrected pSMR. Due to LD, multiple SNPs may be associated with the expression of a gene, and some SNPs are associated with the expression of more than one gene. Since the aim of SMR is to prioritize variants and genes for subsequent studies, a test for heterogeneity excludes regions that may harbor multiple causal loci (pHET < 0.05). SMR analyses were conducted using eQTLGen Consortium, GTEx (11 brain tissues), and CMC data.

We conducted a transcriptome wide association study^65^ using pre-computed expression reference weights for CMC data (5,420 genes with significant cis-SNP heritability) provided with the TWAS/FUSION software. The significance threshold was 0.05/5420.

### DNA looping using Hi-C

Dorsolateral prefrontal cortex (Brodmann area 9) was dissected from postmortem samples from three adults of European ancestry (Dr Craig Stockmeier, University of Mississippi Medical Center). Cerebrum from three fetal brains were obtained from the NIH NeuroBiobank (URLs; gestation age 17-19 weeks, African ancestry). Samples were dry homogenized to a fine powder using a liquid nitrogen-cooled mortar and pestle.

We used “easy Hi-C” (in preparation) to assess DNA looping interactions. Pulverized tissue (~150 mg) was crosslinked with formaldehyde (1% final concentration) and the reaction quenched using glycine (150 mM). Samples were then lysed, Dounce homogenized, and digested using *HindIII*. This was followed by in situ ligation. Samples were cross-linked with proteinase K and purified using phenol-chloroform. DNA was then digested with *DpnII* followed by purification using PCRClean DX beads (Aline Biosciences). The DNA products were self-ligated overnight at 16° using T4 DNA ligase. Self-ligated DNA waw purified with phenol-chloroform, digested with lambda exonuclease, and purified using PCRClean DX beads. For DNA circle re-linearization, bead-bound DNA was eluted and digested with *HindIII* and purified using PCRClean. Bead-bound DNA was eluted in 50ul nuclease free water.

Re-linearized DNA (~50ng) was used for library generation (Illumina TruSeq protocol). Briefly, the DNA was end-repaired using End-it kit (Epicentre), A tailed with Klenow fragment (3′–5′ exo–; NEB), and purified with PCRClean DX beads. The 4ul DNA product was mixed with 5ul of 2X quick ligase buffer, 1ul of 1:10 diluted annealed adapter and 0.5ul of Quick DNA T4 ligase (NEB). The ligation was done by incubating at room temperature for 15 minutes. DNA was purified using DX beads. Elution was done in 14ul nuclease free water. To deep-sequence easy Hi-C libraries, we used custom TruSeq adapter in which the index is replaced by 6 base random sequence. Libraries were then PCR amplified and deeply sequenced (4-5 lanes per sample, around 1 billion reads per sample) using Illumina HiSeq4000 (2x50bp).

Because nearly all mappable reads start with the *HindIII* sequence AGCTT, we trimmed the first 5 bases from every read and added the 6-base sequence AAGCTT to the 5’ of all reads. These read were then aligned to the human reference genome (hg19) using Bowtie. After mapping, we kept reads where both ends were exactly at *HindIII* cutting sites. PCR duplicates were removed. Of these *HindIII* pairs, we split reads into three classes based on their strand orientations (“same-strand”, “inward”, or “outward”). For cis-reads the only type of invalid cis-pairs are self-circles with two ends within the same *HindIII* fragment facing each other. We computed the total number of real cis-contact as twice the number of valid “same-strand” pairs. Reads from undigested *HindIII* sites are back-to-back read pairs next to the same *HindIII* sites facing away from each other.

### Gene-wise and pathway analysis

Our approach was guided by rigorous method comparisons conducted by PGC members.^70,114^ *P*-values quantifying the degree of association of genes and gene sets with MDD were generated using MAGMA (v1.06).^115^ MAGMA uses Brown’s method to combine SNP p-values and account for LD. We used ENSEMBL gene models for 19,079 genes giving a Bonferroni corrected *P*-value threshold of 2.6×10^−6^. Gene set *P*-values were obtained using a competitive analysis that tests whether genes in a gene set are more strongly associated with the phenotype than other gene sets. We used European-ancestry subjects from 1,000 Genomes Project (Phase 3 v5a, MAF ≥ 0.01)^101^ for the LD reference. The gene window used was 35 kb upstream and 10 kb downstream to include regulatory elements.

Gene sets were from two main sources. First, we included gene sets previously shown to be important for psychiatric disorders (71 gene sets; e.g., FMRP binding partners, *de novo* mutations, GWAS top SNPs, ion channels).^72,116,117^ Second, we included gene sets from MSigDB (v5.2)^118^ which includes canonical pathways and Gene Ontology gene sets. Canonical pathways were curated from BioCarta, KEGG, Matrisome, Pathway Interaction Database, Reactome, SigmaAldrich, Signaling Gateway, Signal Transduction KE, and SuperArray. Pathways containing between 10-10K genes were included.

To evaluate gene sets related to antidepressants, gene-sets were extracted from the Drug-Gene Interaction database (DGIdb v.2.0)^119^ and the Psychoactive Drug Screening Program Ki DB^120^ downloaded in June 2016. The association of 3,885 drug gene-sets with MDD was estimated using MAGMA (v1.6). The drug gene-sets were ordered by p-value, and the Wilcoxon-Mann-Whitney test was used to assess whether the 42 antidepressant gene-sets in the dataset (ATC code N06A in the Anatomical Therapeutic Chemical Classification System) had a higher ranking than expected by chance.

One issue is that some gene sets contain overlapping genes, and these may reflect largely overlapping results. The pathway map was constructed using the kernel generative topographic mapping algorithm (k-GTM) as described by Olier et al. GTM is a probabilistic alternative to Kohonen maps: the kernel variant is used when the input is a similarity matrix. The GTM and k-GTM algorithms are implemented in GTMapTool (URLs). We used the Jaccard similarity matrix of FDR-significant pathways as input for the algorithm, where each pathway is encoded by a vector of binary values representing the presence (1) or absence (0) of a gene. Parameters for the k-GTM algorithm are the square root of the number of grid points (k), the square root of the number of RBF functions (m), the regularization coefficient (l), the RBF width factor (w), and the number of feature space dimensions for the kernel algorithm (b). We set k=square root of the number of pathways, m=square root of k, l=1 (default), w=1 (default), and b=the number of principal components explaining 99.5% of the variance in the kernel matrix. The output of the program is a set of coordinates representing the average positions of pathways on a 2D map. The x and y axes represent the dimensions of a 2D latent space. The pathway coordinates and corresponding MAGMA *P*-values were used to build the pathway activity landscape using the kriging interpolation algorithm implemented in the R gstat package.

### Mendelian randomization (MR).^121^

We used MR to investigate the relationships between MDD and correlated traits. Epidemiological studies show that MDD is associated with environmental and life event risk factors as well as multiple diseases, yet it remains unclear whether such trait outcomes are causes or consequences of MDD (or prodromal MDD). Genetic variants are present from birth, and hence are far less likely to be confounded with environmental factors than in epidemiological studies.

We conducted bi-directional MR analysis for four traits: years of education (EDY)^76^, body mass index (BMI)^27^, coronary artery disease (CAD)^77^, and schizophrenia (SCZ)^22^. Briefly, we denote z as a genetic variant (i.e., a SNP) that is significantly associated with x, an exposure or putative causal trait for y (the disease/trait outcome). The effect size of x on y can be estimated using a two-step least squares (2SLS)^122^ approach: 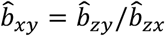, where 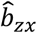 is the estimated effect size for the SNP-trait association the exposure trait, and 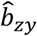 is the effect size estimated for the same SNP in the GWAS of the outcome trait.

Since SNP-trait effect sizes are typically small, power is increased by using multiple associated SNPs which allows simultaneous investigation of pleiotropy driving the epidemiologically observed trait associations. Causality of the exposure trait for the outcome trait implies a consistent relationship between the SNP association effect sizes of the exposure associated SNPs in the outcome trait.

We used generalized summary statistics-based MR (GSMR) (Zhu et al., submitted) to estimate 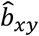 and its standard error from multiple SNPs associated with the exposure trait at a genome-wide significance level. We conducted bi-directional GSMR analyses for each pair of traits, and report results after excluding SNPs that fail the HEIDI-outlier heterogeneity test (which is more conservative than excluding SNPs that have an outlying association likely driven by locus-specific pleiotropy). GSMR is more powerful than inverse-weighted MR (IVW-MR) and MR-Egger because it takes account of the sampling variation of both 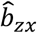 and 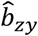. GSMR also accounts for residual LD between the clumped SNPs. For comparison, we also conducted IVW-MR and MR-Egger analyses.^123^

### Trans-ancestry

Common genetic risk variants for complex biomedical conditions are likely to be shared across ancestries.^124,125^ However, lower *r_g_* have been reported likely reflecting different LD patterns by ancestry. For example, European-Chinese *r_g_* estimates were below one for ADHD (0.39, SE 0.15),^126^ rheumatoid arthritis (0.46, SE 0.06),^127^ and type 2 diabetes (0.62, SE 0.09),^127^ and reflect population differences in LD and population-specific causal variants.

The Han Chinese CONVERGE study^17^ included clinically ascertained females with severe, recurrent MDD, and is the largest non-European MDD GWA to date. Neither of the two genome-wide significant loci in CONVERGE had SNP findings ±250 kb with *P* < 1×10^−6^ in the full European results. We used LDSC with an ancestry-specific LD reference for within ancestry estimation, and POPCORN^127^ for trans-ancestry estimation. In the CONVERGE sample, 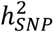was reported as 20-29%.^128^ Its *r_g_* with the seven European MDD cohorts was 0.33 (SE 0.03).^129^ For comparison, *r_g_* for CONVERGE with European results for schizophrenia was 0.34 (SE 0.05) and 0.45 (SE 0.07) for bipolar disorder. The weighted mean *r_g_* between the CONVERGE cohort with the seven anchor and expanded cohorts using was 0.31 (SE 0.03). These *r_g_* estimates should be interpreted in light of the estimates of *r_g_* within European MDD cohorts which are variable (***Table S4***).

### Genome build

All genomic coordinates are given in NCBI Build 37/UCSC hg19.

### Availability of results

The PGC’s policy is to make genome-wide summary results public. Summary statistics for a combined meta-analysis of the anchor cohort samples with five of the six expanded samples (deCODE, Generation Scotland, GERA, iPSYCH, and UK Biobank) are available on the PGC web site (URLs). Results for 10,000 SNPs for all seven cohorts are also available on the PGC web site.

GWA summary statistics for the sixth expanded cohort (23andMe, Inc.) must be obtained separately. Summary statistics for the 23andMe dataset can be obtained by qualified researchers under an agreement with 23andMe that protects the privacy of the 23andMe participants. Please contact David Hinds for more information and to apply to access the data. Researchers who have the 23andMe summary statistics can readily recreate our results by meta-analyzing the six cohort results file with the Hyde et al. results file from 23andMe.^19^

<u>Availability of genotype data</u> for the anchor cohorts is described in ***Table S14***. For the expanded cohorts, interested users should contact the lead PIs of these cohorts (which are separate from the PGC).

## URLs

1000 Genomes Project multi-ancestry imputation panel, https://mathgen.stats.ox.ac.uk/impute/data_download_1000G_phase1_integrated.html

23andMe privacy policy https://www.23andme.com/en-eu/about/privacy

Bedtools, https://bedtools.readthedocs.io

Genotype-based checksums for relatedness determination, http://www.broadinstitute.org/~sripke/share_links/checksums_download

GTEx, http://www.gtexportal.org/home/datasets

GTMapTool, http://infochim.u-strasbg.fr/mobyle-cgi/portal.py#forms::gtmaptool

LD-Hub, http://ldsc.broadinstitute.org

MDD summary results are available on the PGC website, https://pgc.unc.edu

NIH NeuroBiobank, https://neurobiobank.nih.gov

PGC “ricopili” GWA pipeline, https://github.com/Nealelab/ricopili

UK Biobank, http://www.ukbiobank.ac.uk

## Author Contributions

### Writing group

G. Breen, A. D. Børglum, D. F. Levinson, C. M. Lewis, S. Ripke, P. F. Sullivan, N. R. Wray.

### PGC MDD PI group

V. Arolt, B. T. Baune, K. Berger, D. I. Boomsma, G. Breen, A. D. Børglum, S. Cichon, U. Dannlowski, J. R. DePaulo, E. Domenici, K. Domschke, T. Esko, E. d. Geus, H. J. Grabe, S. P. Hamilton, C. Hayward, A. C. Heath, D. M. Hougaard, K. S. Kendler, S. Kloiber, D. F. Levinson, C. M. Lewis, G. Lewis, Q. S. Li, S. Lucae, P. A. Madden, P. K. Magnusson, N. G. Martin, A. M. McIntosh, A. Metspalu, O. Mors, P. B. Mortensen, B. Müller-Myhsok, M. Nordentoft, M. M. Nöthen, M. C. O’Donovan, S. A. Paciga, N. L. Pedersen, B. W. Penninx, R. H. Perlis, D. J. Porteous, J. B. Potash, M. Preisig, M. Rietschel, C. Schaefer, T. G. Schulze, J. W. Smoller, K. Stefansson, P. F. Sullivan, H. Tiemeier, R. Uher, H. Völzke, M. M. Weissman, T. Werge, A. R. Winslow, N. R. Wray.

### Bioinformatics

23andMe Research Team, M. J. Adams, S. V. d. Auwera, G. Breen, J. Bryois, A. D. Børglum, E. Castelao, J. H. Christensen, T. Clarke, J. R. I. Coleman, L. Colodro-Conde, eQTLGen Consortium, G. E. Crawford, C. A. Crowley, G. Davies, E. M. Derks, T. Esko, A. J. Forstner, H. A. Gaspar, P. Giusti-Rodríguez, J. Grove, L. S. Hall, T. F. Hansen, C. Hayward, M. Hu, R. Jansen, F. Jin, Z. Kutalik, Q. S. Li, Y. Li, P. A. Lind, X. Liu, L. Lu, D. J. MacIntyre, S. E. Medland, E. Mihailov, Y. Milaneschi, J. N. Painter, B. W. Penninx, W. J. Peyrot, G. Pistis, P. Qvist, L. Shen, S. I. Shyn, C. A. Stockmeier, P. F. Sullivan, K. E. Tansey, A. Teumer, P. A. Thomson, A. G. Uitterlinden, Y. Wang, S. M. Weinsheimer, N. R. Wray, H. S. Xi.

### Clinical

E. Agerbo, T. M. Air, V. Arolt, B. T. Baune, A. T. F. Beekman, K. Berger, E. B. Binder, D. H. R. Blackwood, H. N. Buttenschøn, A. D. Børglum, N. Craddock, U. Dannlowski, J. R. DePaulo, N. Direk, K. Domschke, M. Gill, F. S. Goes, H. J. Grabe, A. C. Heath, A. M. v. Hemert, I. B. Hickie, M. Ising, S. Kloiber, J. Krogh, D. F. Levinson, S. Lucae, D. J. MacIntyre, D. F. MacKinnon, P. A. Madden, W. Maier, N. G. Martin, P. McGrath, P. McGuffin, A. M. McIntosh, A. Metspalu, C. M. Middeldorp, S. S. Mirza, F. M. Mondimore, O. Mors, P. B. Mortensen, D. R. Nyholt, H. Oskarsson, M. J. Owen, C. B. Pedersen, M. G. Pedersen, J. B. Potash, J. A. Quiroz, J. P. Rice, M. Rietschel, C. Schaefer, R. Schoevers, E. Sigurdsson, G. C. B. Sinnamon, D. J. Smith, F. Streit, J. Strohmaier, D. Umbricht, M. M. Weissman, J. Wellmann, T. Werge, G. Willemsen.

### Genomic assays

G. Breen, H. N. Buttenschøn, J. Bybjerg-Grauholm, M. Bækvad-Hansen, A. D. Børglum, S. Cichon, T. Clarke, F. Degenhardt, A. J. Forstner, S. P. Hamilton, C. S. Hansen, A. C. Heath, P. Hoffmann, G. Homuth, C. Horn, J. A. Knowles, P. A. Madden, L. Milani, G. W. Montgomery, M. Nauck, M. M. Nöthen, M. Rietschel, M. Rivera, E. C. Schulte, T. G. Schulze, S. I. Shyn, H. Stefansson, F. Streit, T. E. Thorgeirsson, J. Treutlein, A. G. Uitterlinden, S. H. Witt, N. R. Wray.

### Obtained funding for primary MDD samples

B. T. Baune, K. Berger, D. H. R. Blackwood, D. I. Boomsma, G. Breen, H. N. Buttenschøn, A. D. Børglum, S. Cichon, J. R. DePaulo, I. J. Deary, E. Domenici, T. C. Eley, T. Esko, H. J. Grabe, S. P. Hamilton, A. C. Heath, D. M. Hougaard, I. S. Kohane, D. F. Levinson, C. M. Lewis, G. Lewis, Q. S. Li, S. Lucae, P. A. Madden, W. Maier, N. G. Martin, P. McGuffin, A. M. McIntosh, A. Metspalu, G. W. Montgomery, O. Mors, P. B. Mortensen, M. Nordentoft, D. R. Nyholt, M. M. Nöthen, P. F. O’Reilly, B. W. Penninx, D. J. Porteous, J. B. Potash, M. Preisig, M. Rietschel, C. Schaefer, T. G. Schulze, G. C. B. Sinnamon, J. H. Smit, D. J. Smith, H. Stefansson, K. Stefansson, P. F. Sullivan, T. E. Thorgeirsson, H. Tiemeier, A. G. Uitterlinden, H. Völzke, M. M. Weissman, T. Werge, N. R. Wray.

### Statistical analysis

23andMe Research Team, A. Abdellaoui, M. J. Adams, T. F. M. Andlauer, S. V. d. Auwera, S. Bacanu, K. Berger, T. B. Bigdeli, G. Breen, E. M. Byrne, A. D. Børglum, N. Cai, T. Clarke, J. R. I. Coleman, B. Couvy-Duchesne, H. S. Dashti, G. Davies, N. Direk, C. V. Dolan, E. C. Dunn, N. Eriksson, V. Escott-Price, T. Esko, H. K. Finucane, J. Frank, H. A. Gaspar, S. D. Gordon, J. Grove, L. S. Hall, C. Hayward, A. C. Heath, S. Herms, D. A. Hinds, J. Hottenga, C. L. Hyde, M. Ising, E. Jorgenson, F. F. H. Kiadeh, J. Kraft, W. W. Kretzschmar, Z. Kutalik, J. M. Lane, C. M. Lewis, Q. S. Li, Y. Li, D. J. MacIntyre, P. A. Madden, R. M. Maier, J. Marchini, M. Mattheisen, H. Mbarek, A. M. McIntosh, S. E. Medland, D. Mehta, E. Mihailov, Y. Milaneschi, S. S. Mirza, S. Mostafavi, N. Mullins, B. Müller-Myhsok, B. Ng, M. G. Nivard, D. R. Nyholt, P. F. O’Reilly, R. E. Peterson, E. Pettersson, W. J. Peyrot, G. Pistis, D. Posthuma, S. M. Purcell, B. P. Riley, S. Ripke, M. Rivera, R. Saxena, C. Schaefer, L. Shen, J. Shi, S. I. Shyn, H. Stefansson, S. Steinberg, P. F. Sullivan, K. E. Tansey, H. Teismann, A. Teumer, W. Thompson, P. A. Thomson, T. E. Thorgeirsson, C. Tian, M. Traylor, V. Trubetskoy, M. Trzaskowski, A. Viktorin, P. M. Visscher, Y. Wang, B. T. Webb, J. Wellmann, T. Werge, N. R. Wray, Y. Wu, J. Yang, F. Zhang.

## Acknowledgements

PGC: We are deeply indebted to the investigators who comprise the PGC, and to the hundreds of thousands of subjects who have shared their life experiences with PGC investigators. Statistical analyses were carried out on the NL Genetic Cluster Computer (http://www.geneticcluster.org) hosted by SURFsara.

EDINBURGH: Genotyping was conducted at the Genetics Core Laboratory at the Clinical Research Facility (University of Edinburgh). GenScot: We are grateful to all the families who took part, the general practitioners and the Scottish School of Primary Care for their help in recruiting them, and the whole Generation Scotland team, which includes interviewers, computer and laboratory technicians, clerical workers, research scientists, volunteers, managers, receptionists, healthcare assistants and nurses. Genotyping was conducted at the Genetics Core Laboratory at the Clinical Research Facility (University of Edinburgh). GSK_MUNICH: We thank all participants in the GSK-Munich study. We thank numerous people at GSK and Max-Planck Institute, BKH Augsburg and Klinikum Ingolstadt in Germany who contributed to this project. JANSSEN: Funded by Janssen Research & Development, LLC. We are grateful to the study volunteers for participating in the research studies and to the clinicians and support staff for enabling patient recruitment and blood sample collection. We thank the staff in the former Neuroscience Biomarkers of Janssen Research & Development for laboratory and operational support (e.g., biobanking, processing, plating, and sample de-identification), and to the staff at Illumina for genotyping Janssen DNA samples. MARS: This work was funded by the Max Planck Society, by the Max Planck Excellence Foundation, and by a grant from the German Federal Ministry for Education and Research (BMBF) in the National Genome Research Network framework (NGFN2 and NGFN-Plus, FKZ 01GS0481), and by the BMBF Program FKZ 01ES0811. We acknowledge all study participants. We thank numerous people at Max-Planck Institute, and all study sites in Germany and Switzerland who contributed to this project. Controls were from the Dortmund Health Study which was supported by the German Migraine & Headache Society, and by unrestricted grants to the University of Münster from Almirall, Astra Zeneca, Berlin Chemie, Boehringer, Boots Health Care, Glaxo-Smith-Kline, Janssen Cilag, McNeil Pharma, MSD Sharp & Dohme, and Pfizer. Blood collection was funded by the Institute of Epidemiology and Social Medicine, University of Münster. Genotyping was supported by the German Ministry of Research and Education (BMBF grant 01ER0816). PsyColaus: PsyCoLaus/CoLaus received additional support from research grants from GlaxoSmithKline and the Faculty of Biology and Medicine of Lausanne. QIMR: We thank the twins and their families for their willing participation in our studies. RADIANT: This report represents independent research funded by the National Institute for Health Research (NIHR) Biomedical Research Centre at South London and Maudsley NHS Foundation Trust, and King’s College London. The views expressed are those of the authors and not necessarily those of the NHS, the NIHR, or the Department of Health. Rotterdam Study: The Rotterdam Study is also funded by Erasmus Medical Center and Erasmus University. SHIP-LEGEND/TREND: SHIP is part of the Community Medicine Research net of the University of Greifswald which is funded by the Federal Ministry of Education and Research (grants 01ZZ9603, 01ZZ0103, and 01ZZ0403), the Ministry of Cultural Affairs, and the Social Ministry of the Federal State of Mecklenburg-West Pomerania. Genotyping in SHIP was funded by Siemens Healthineers and the Federal State of Mecklenburg-West Pomerania. Genotyping in SHIP-TREND-0 was supported by the Federal Ministry of Education and Research (grant 03ZIK012). STAR*D: The authors appreciate the efforts of the STAR*D investigator team for acquiring, compiling, and sharing the STAR*D clinical data set. TwinGene: thanks the Karolinska Institutet for infrastructural support of the Swedish Twin Registry. 23andME: We thank the 23andMe research participants included in the analysis, all of whom provided informed consent and participated in the research online according to a human subjects protocol approved by an external AAHRPP-accredited institutional review board (Ehical & Independent Review Services), and the employees of 23andMe for making this work possible. 23andMe acknowledges the invaluable contributions of Michelle Agee, Babak Alipanahi, Adam Auton, Robert K. Bell, Katarzyna Bryc, Sarah L. Elson, Pierre Fontanillas, Nicholas A. Furlotte, David A. Hinds, Bethann S. Hromatka, Karen E. Huber, Aaron Kleinman, Nadia K. Litterman, Matthew H. McIntyre, Joanna L. Mountain, Carrie A.M. Northover, Steven J. Pitts, J. Fah Sathirapongsasuti, Olga V. Sazonova, Janie F. Shelton, Suyash Shringarpure, Chao Tian, Joyce Y. Tung, Vladimir Vacic, and Catherine H. Wilson. deCODE: The authors are thankful to the participants and staff at the Patient Recruitment Center. GERA: Participants in the Genetic Epidemiology Research on Adult Health and Aging Study are part of the Kaiser Permanente Research Program on Genes, Environment, and Health, supported by the Wayne and Gladys Valley Foundation, The Ellison Medical Foundation, the Robert Wood Johnson Foundation, and the Kaiser Permanente Regional and National Community Benefit Programs. iPSYCH: The iPSYCH (The Lundbeck Foundation Initiative for Integrative Psychiatric Research) team acknowledges funding from The Lundbeck Foundation (grant no R102-A9118 and R155-2014-1724), the Stanley Medical Research Institute, the European Research Council (project no: 294838), the Novo Nordisk Foundation for supporting the Danish National Biobank resource, and grants from Aarhus and Copenhagen Universities and University Hospitals, including support to the iSEQ Center, the GenomeDK HPC facility, and the CIRRAU Center. UK Bioband: this research has been conducted using the UK Biobank Resource (URLs), including applications #4844 and #6818. Finally, we thank the members of the eQTLGen Consortium for allowing us to use their very large eQTL database ahead of publication. Its members are listed in ***Table S15***.

## Funding sources

The table below lists the funding that supported the primary studies analyzed in the paper.

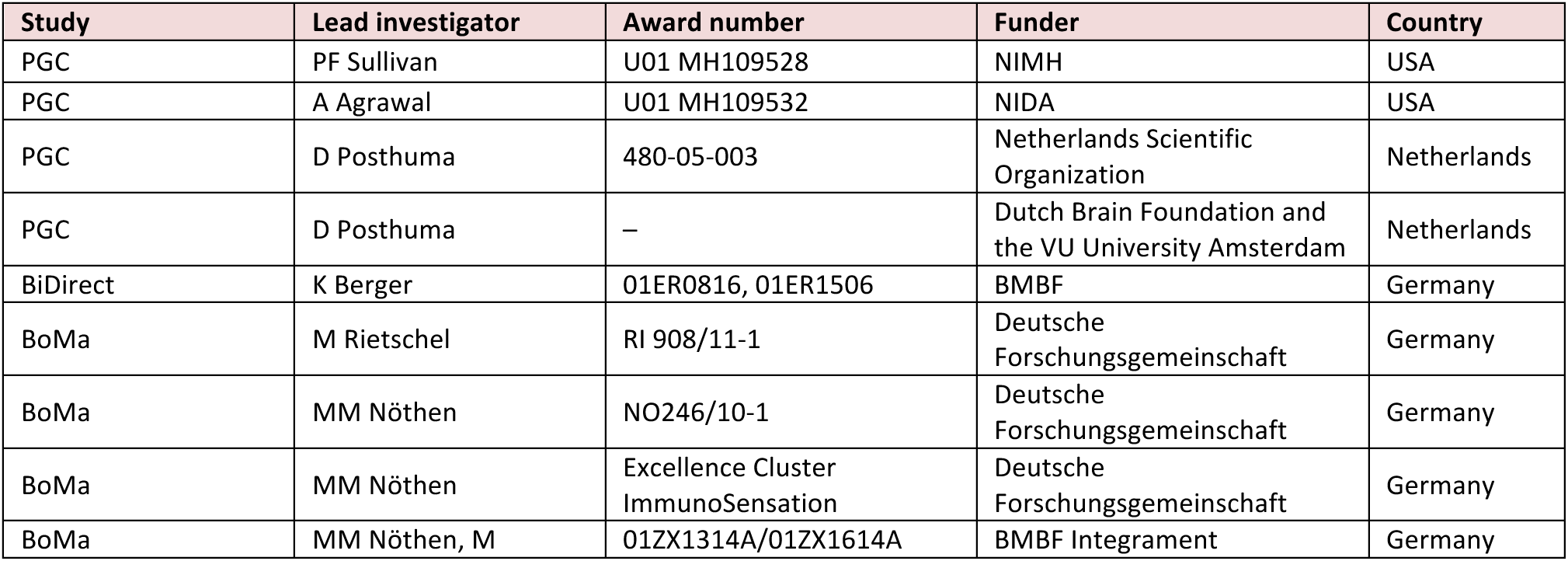
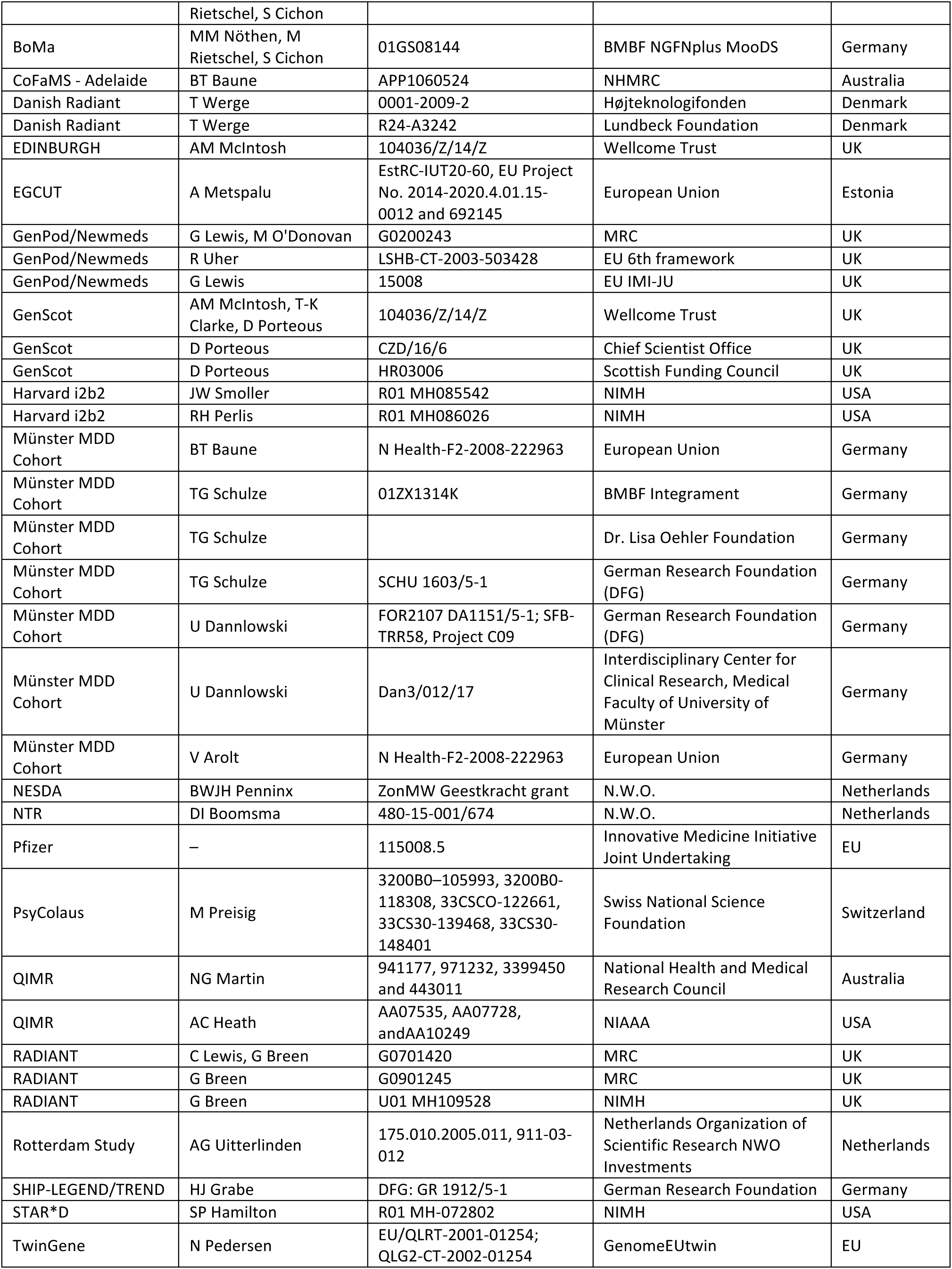
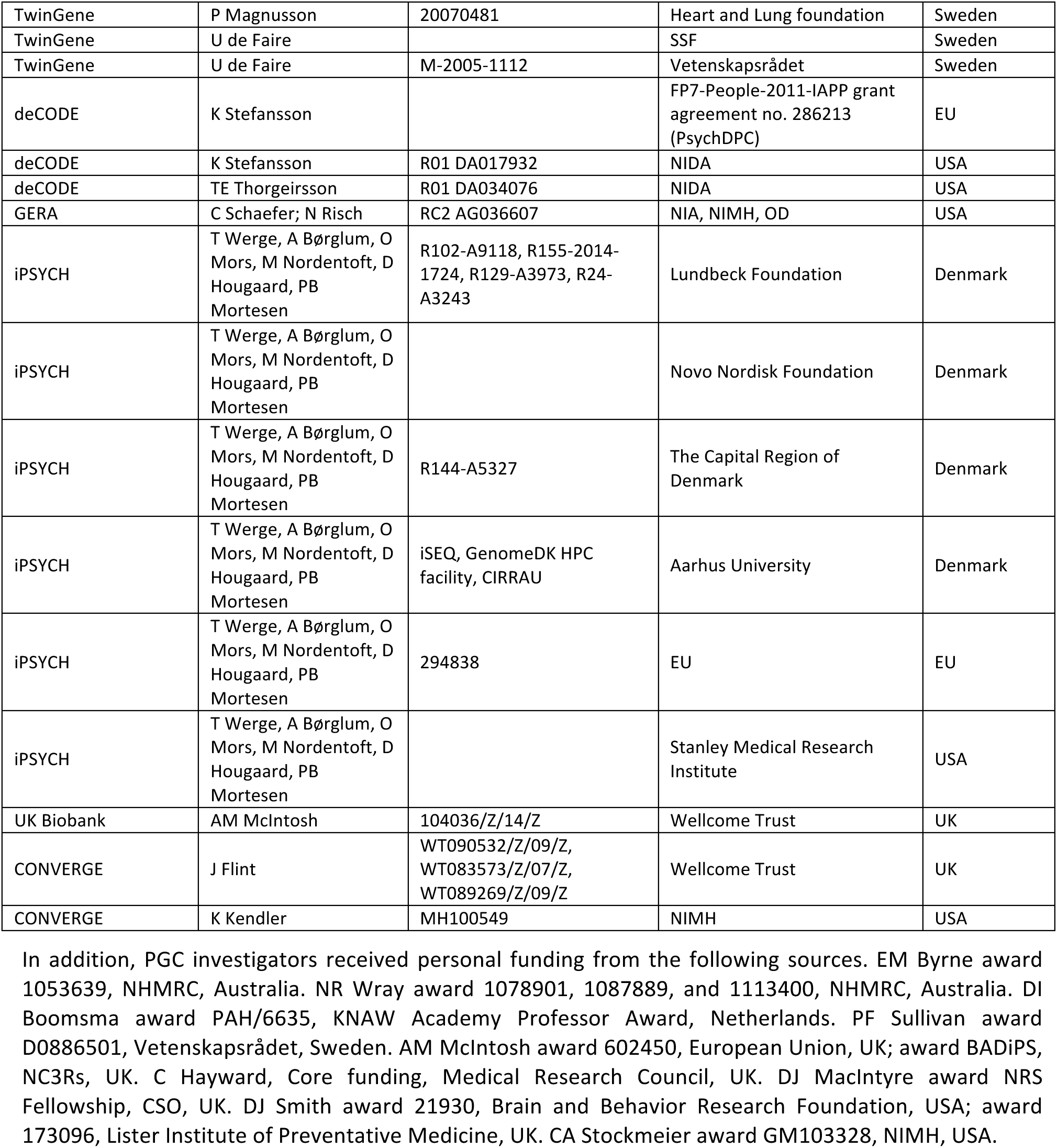

---

**Figure S1:** Leave-one-out GRS analyses of the anchor cohort. (a) Per sample R^2^ at varying significance thresholds. A all samples in the anchor cohort (except one) yielded significant differences in case-control distributions of GRS. Across all samples in the anchor cohort, GRS explained 1.9% of variance in liability. (b) Relation between the number of cases and R^2^, showing the expected positive correlation.

**Figure S2:** Regional association plots of genomic regions identified from SMR analysis of MDD GWA and eQTL results. SMR analysis helps to prioritize specific genes in a region of association for follow-up functional studies. Figures appear in the same order as the results reported in ***Table S9***. In the top plot, grey dots represent the MDD GWA *P*-values, diamonds show *P*-values for probes from the SMR test, and triangles are probes without a cis-eQTL (at *P_eQTL_* < 5e-8). Genes that pass SMR and heterogeneity tests (designed to remove loci with more than one causal association) are highlighted in red. The eQTL *P*-values of SNPs are shown in the bottom plot.

**Figure S3:** Circular plots to illustrate DNA-DNA loops. From the outside, the tracks show hg19 coordinates in Mb, the positions of significant MDD associations (−log_10_(P), outward is more significant), the names and positons of GENCODE genes, and the arc show significant DNA-DNA loops (q < 1e-4) from Hi-C on adult cortex (green) and fetal frontal cortex (blue). (a) chr1:71.5-74.1 Mb suggesting that the two statistically independent associations in the region both implicate NEGR1. (b) The MDD association in RERE, in contrast, coincides with many DNA-DNA loops and may suggest that this region contains super-enhancer elements.

**Figure S4:** Graphs depicting the SNP instruments used in Mendelian randomization analyses. Table S13 shows the parameter estimates and significance, and these graphs show scatterplots of the instruments for MDD and (a) BMI, (b) years of education, (c) coronary artery disease, and (d) schizophrenia.

